# *Staphylococcus sciuri* C2865 from a distinct subspecies cluster as reservoir of the novel transferable trimethoprim resistance gene, *dfrE,* and adaptation driving mobile elements

**DOI:** 10.1101/2020.09.30.320143

**Authors:** Elena Gómez-Sanz, Jose Manuel Haro-Moreno, Slade O. Jensen, Juan José Roda-García, Mario López-Pérez

## Abstract

Four methicillin-resistant *Staphylococcus sciuri* (MRSS) strains isolated from stranded dogs showed trimethoprim (TMP) resistance, while all staphylococcal TMP resistant dihydrofolate reductase genes (*dfr*) were negative. An in-depth whole-genome-sequencing approach on strain C2865 was followed for *resistome* and *mobilome* profiling, and for comparative genomics with *S. sciuri* group available genomes. Lack of species host tropism was observed, with MRSS C2865 placed at a separate sub-branch within *S. sciuri* species, close to the average nucleotide identity to be considered a different species (95-96%). *S. sciuri* proved a pronounced accessory genome (73% of genes), while MRSS C2865 distinctively harboured the highest total gene number and highest number of unique genes, with 75% associated to the recognised *mobilome*. A novel multidrug resistance mosaic plasmid (pUR2865-34) with several adaptive, mobilization (*oriT* mimic) and segregational stability (Type Ib *par* system) traits and two small single resistance plasmids were identified. Plasmid pUR2865-34 enclosed a novel staphylococcal TMP resistance gene, named *dfrE*, which shared highest identity with *dfr* of soil-related *Paenibacillus anaericanus* (68%). DfrE conferred high-level TMP resistance in *S. aureus* and *Escherichia coli*. Database searches revealed that *dfrE* was formerly denoted (*dfr_like*) in an *Exiguobacterium* spp. from a fish-farm sediment and that was present but unnoticed in several staphylococcal and one macrococcal genomes with different epidemiological backgrounds. Novel chromosomal site-specific mobile islands with resourceful traits were identified, including a multidrug-resistant SCC*<u>mec</u>* cassette lacking cassette chromosome recombinase (Ccr) genes, a staphylococcal pathogenicity island of the SaPI4 family, and three unrelated *siphoviridae* prophages, two of which enclosed recombinases with the conserved Ccr-motif. We reveal a novel staphylococcal TMP resistance *dfrE* gene already present in diverse bacterial backgrounds. We confirm the ubiquity, high genome plasticity and low host tropism of *S. sciuri* highlighting its role as a resourceful reservoir for evolutionary novel features contributing to its extraordinary versatility and adaptability.

**Author summary:** *Staphylococcus* spp. are ubiquitous bacteria present in diverse ecological niches, including humans, animals and the environment. They are clinically relevant opportunistic pathogens and are notorious for their ability to acquire antimicrobial resistance (AMR) and virulence properties, resulting in a significant impact for Public Health. Mobile genetic elements (MGEs) play a central role in this adaptation process and are a means to transfer genetic information across bacterial species. *Staphylococcus sciuri* represents one of the most ancestral species in the genus and has been suggested a reservoir for AMR genes. Here, following a refined whole genome sequencing approach we determined the entire genome of an animal and environment-associated multidrug resistant (MDR) *S. sciuri* strain uncovering a novel acquired staphylococcal TMP resistance gene already spread among different bacterial species from different epidemiological backgrounds. We also reveal several additional MGEs, including a novel MDR mobilizable plasmid that encloses several adaptive and stabilization features, and novel mobilizable chromosomal islands with resourceful traits, including three unrelated prophages. Together with comparative genomics, we confirm the ubiquity, high intraspecies heterogenicity, genome plasticity and low host tropism of this species, highlighting its role as resourceful reservoir for evolutionary novel features contributing to its extraordinary versatility and adaptability.

## Introduction

*Staphylococcus* spp. are ubiquitous bacteria present in diverse ecological niches. They are members of the natural microbiota of the skin and mucosa of humans, other mammals and birds. They are also opportunistic pathogens, responsible for mild to life-threatening infections, such as skin and soft tissue infections, food poisoning, endocarditis, and sepsis [1]. Coagulase negative staphylococci (CoNS), the major group within the genus, now represent one of the major nosocomial pathogens [2]. Within this cluster, *Staphylococus sciuri* species subgroup includes five species (*S. sciuri*, *Staphylococus lentus*, *Staphylococus vitulinus*, *Staphylococus fleurettii* and *Staphylococus stepanovicii*) that are most often present as commensal animal-associated bacteria [3]. *S. sciuri*, taxonomically considered the most primitive species among staphylococci, has been shown to present high intraspecies heterogeneity. However, its traditional classification in three distinct subspecies is currently rejected [4]. The ubiquitous presence of *S. sciuri* represents a continuous source for contamination, colonization and infection in animals and humans from different niches, including dust and hospital surfaces [5–13]. Direct animal contact or food products of animal origin may be extra sources for humans [3, 14, 15].

*S. sciuri* is a natural reservoir of the ancestral *mecA* gene, which evolution generated β-lactam resistance [16–18]. This species is considered a source of its horizontal gene transfer (HGT), via the Staphylococcal Cassette Chromosome (SCC*mec*) mobile element, into other staphylococcal species [18]. *S. sciuri* is frequently multidrug resistant (MDR) and novel clinically relevant AMR genes in staphylococci have been first detected in this species. This includes the multidrug resistance (PhLOPS phenotype) *cfr* gene [19], the macrolide/lincosamide/streptogramin B (MLSB) resistance *erm*(33) [20], the lincosamide/streptogramin A resistance *sal*(A) [21], the oxazolidinone/phenicol resistance *optrA* [22], or the co-existence of plasmid located *cfr-optrA* [22] and the β-lactam resistance *mecA-mecC* in hybrid SCC*mec* elements [23, 24]. *S. sciuri* has also been found to carry several virulence factors, including those implicated in biofilm formation, although potential genes associated to this phenotype have not been specified [3, 25].

Mobile genetic elements (MGEs) play a key role in intra- and interspecies HGT of AMR and virulence determinants. Although some MGEs have been identified in *S. sciuri*, their “mobilome” (pool of genes within MGEs) and diversity remain largely unknown. In particular, despite staphylococcal phages are considered ubiquitous in this genus, and their proven contribution in pathogenesis, virulence and genome plasticity, phages of *S. sciuri* have only been reported twice [26, 27]. Former reports indicate that staphylococcal phages also contribute to the horizontal spread of AMR elements [28–32], including mobilization of the SCC*mec* element [27, 30, 33, 34].

In staphylococci, resistance to trimethoprim (TMP) is mediated by any of the following acquired dihydrofolate reductases (Dfrs): DfrA (DfrS1), DfrD, DfrG, DfrK and DfrF [35–40], which are TMP insensitive variants of the intrinsic dihydrofolate reductase/s (Dhfr). In this study, a canine methicillin-resistant *S. sciuri* (MRSS) strain C2865 from Nigeria showed TMP resistance, while all staphylococcal *dfr* genes were negative. A sequential whole-genome-sequencing (WGS) approach was conducted aiming to determine not only the genetic basis for TMP resistance, but also the complete “resistome” (pool of AMR determinants) and mobilome profiling and a comparative genomic analyses with former *S. sciuri* species group members. Our work unveils a novel mobilizable staphylococcal TMP resistance gene, *dfrE*, a number of novel mobile elements with resistance and further adaptive and stabilization features and confirms the high genome plasticity of this species. Our results highlight the importance of *S. sciuri* as a reservoir for novel mobilizable AMR determinants and MGEs with adaptive features to other staphylococci.

## Results

### Sequencing approach reasoning and general characteristics of *S. sciuri* C2865 genome

Assembly of trimmed Illumina sequence reads resulted in 341 contigs with a contig sum of 2,937,715 bp. Among these, four different plasmids were apparently detected, pUR2865-1 (2’559 bp), pUR2865-2 (3’830 bp), pUR2865-3 (14’849 bp) and pUR2865-4 (25’259 bp). Plasmids pUR2865-1 and pUR2865-2 were located as single circularized contigs, based on contig boundary redundancy. On the other hand, putative pUR2865-3 and pUR2865-4 had no clear delimitation and PCR assays could not drag conclusive results. In addition, several fragmented chromosomally located mobile elements were detected; however, reiterate attempts to determine their entire structure and delimitation failed. This flexible genome enclosed several ISs, transposases and integrases that were responsible for the assembly disruption. Subsequently, PacBio sequencing selecting a large DNA fragmentation size (15 kb) was pursuit for genome and mobilome completeness, accepting the loss of both small (<4 kb) plasmids. Assembly of trimmed PacBio sequence reads resulted in one single circular chromosomal contig of 2,913,767 bp plus one circularized 40,108-bp plasmid (pUR2865-34), making a sum of 2,953,875 bp (16,160 bp more than Illumina). When including the size of the two small plasmids missed with PacBio (pUR2865-1, pUR2865-2), *S. sciuri* C2865 genome consisted of 2,960,715 bp. Summary of MRSS C2865 sequencing and assembly data generated by Illumina and PacBio is shown in Table 1. General characteristics of MRSS C2865 MGEs detected is displayed in Table 2.

**Table 1.** Sequencing and assembly data comparison of the *S. sciuri* C2865 genome processed with Illumina Miseq and with PacBio RSII.

|  | Parameter | Illumina MiSeq | PacBio RSII |
| --- | --- | --- | --- |
| Sequencing | Number of bp | 662,993,478 | 2,162,733,205 |
|  | Number of reads | 2,774,428 | 103,918 |
|  | Mean read length | 239 | 20,811 |
|  | Average genome coverage | 225x | 730x |
| Assembly | Total assembled sequence (bp) | 2,937,715 | 2,953,875 |
|  | Total assembled contigs | 341 | 2 |
|  | Mean contig size | 8,615 | 1,488,849 |
|  | Maximum contig length | 125,700 | 2,913,767 |
|  | Length of Chromosome sequence (bp) | 2,893,295 | 2,913,767 |
|  | G + C content (%) | 32.7 | 32.5 |
|  | N50 contig length | 37,104 | 2,913,767 |
|  | ORFs | 3,270 | 3,097 |
|  | Gene density (no genes/kb) | 1.13 | 1.06 |
|  | Coding (%) | 91% | 89% |
|  | Median Intergenic Spacer (bp) | 51 | 59 |
|  | Protein coding genes | 3,193 | 3,020 |
|  | Ribosomal RNAs: 16S, 23S, 5S | 19 | 19 |
|  | Transfer RNAs | 58 | 58 |
| Mobile Elements | Pre-identified Insertion Sequences | 22 (15 different) | 78 (18 different) |
|  | Prophage | 2-3 | 3 |
|  | SCC<sub>mec</sub> | 1 | 1 |
|  | Staphylococcal pathogenecity island | 1 | 1 |
|  | Plasmid<sup>a</sup> | 4 | 1 |

<sup>a</sup> Consensus (Illumina + PacBio data) = 3 plasmids. All other definitive values in text were taken from PacBio data.

**Table 2.**
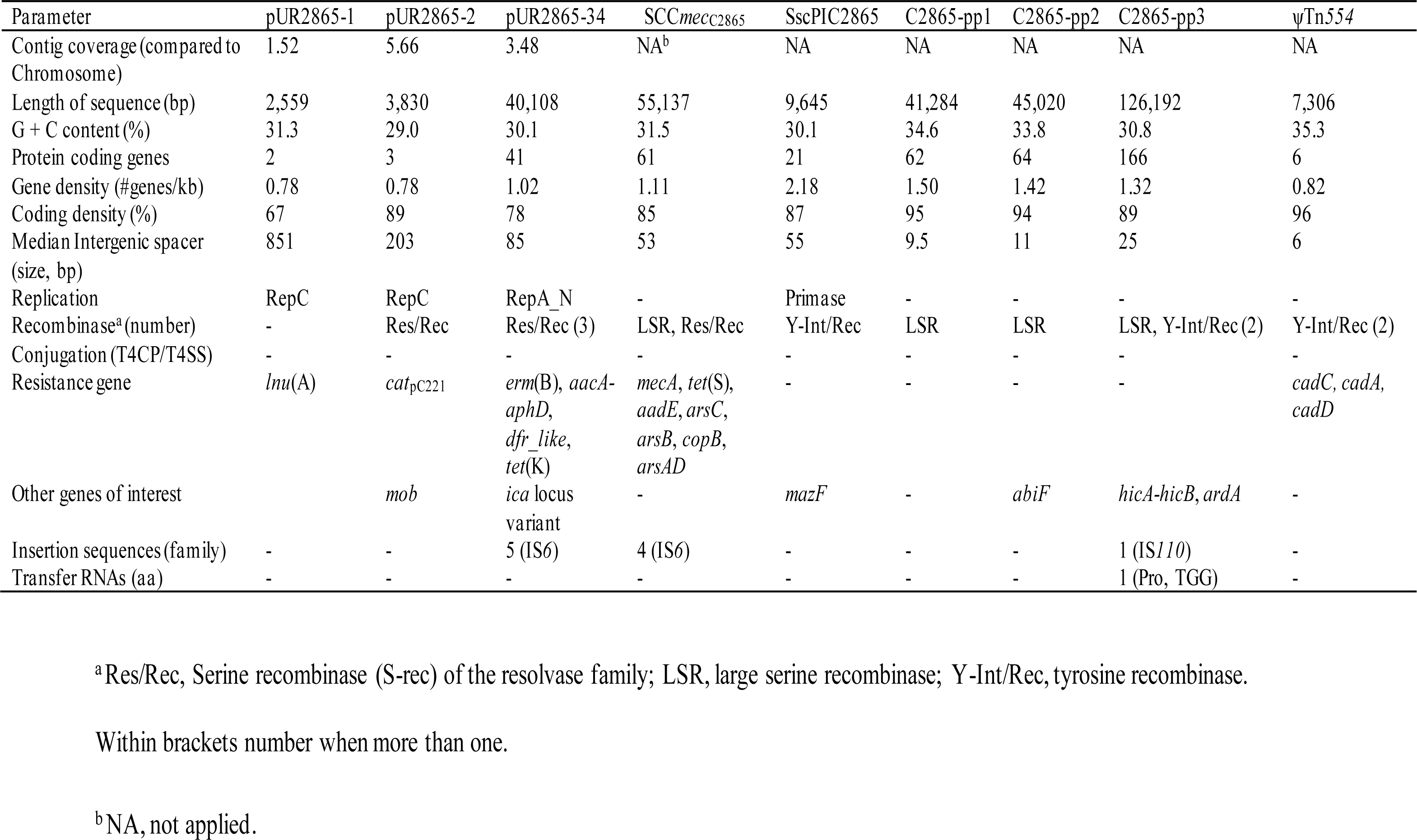
General characteristics of most relevant S. *sciuri* C2865 mobile genetic elements detected as consensus of sequencing data.

Based on the resultant genome-genome alignments using dot-plots of assembled Illumina contigs against the PacBio contigs (reference), 1.9% of the PacBio assembled genome showed no alignment (S1 Fig). Instead, when using the Illumina data as reference, 3.2% of Illumina contigs had no matching (S1 Fig). These unaligned Illumina segments were associated with (i) both small plasmids, (ii) one contig corresponding to plausible contamination and (iii) 166 contigs smaller than 800 bp, many carrying truncated ISs (summing up to 3.1% of Illumina genome). Subsequently, all downstream analyses, except for the two small Illumina-sequenced plasmids, made use of the PacBio data.

A total of 78 ISs were predicted throughout the chromosome of *S. sciuri* strain C2865 using PacBio (data not shown). Among them, 18 were different and belonged to the following families (number of different ISs/total ISs): IS*6* (2/4), IS*200*_IS*605*_ssgr_IS*200* (1/3), IS*256* (2/11), IS*NCY* (2/2), IS*3*_ssgr_IS*150* (6/29), IS*1182* (1/21), IS*21* (1/4), IS*3* (2/2) and IS*110* (1/2). Remarkably, only one IS was validated by the system, due to low percentage of identity with deposited ISs in ISfinder (https://isfinder.biotoul.fr/) implying that most of the identified ISs were new. The plasmid retrieved by PacBio predicted and validated five ISs (two different belonging to IS*6* family). Illumina data produced 22 predicted ISs (15 different) (Table 1), most likely due to the collapse of the repeated regions within the ISs.

### MRSS C2865 belongs to a distinct sub-cluster within *S. sciuri* species and is armed with unique adaptive features

Average nucleotide identity (ANI) of *S. sciuri* C2865 and all *S. sciuri* group genomes deposited in the National Center for Biotechnology Information (NCBI) revealed that *S. sciuri* C2865 shared highest identity with multidrug resistant *S. sciuri* Z8 (99% ANI) isolated from a human skin wound infection in China [41], MDR *S. sciuri* SNUDS-18 (98%), from a duckling with tremor in South Korea [42] and *S. sciuri* CCUG39509 (98%), from a sliced veal leg in Sweden (RefSeq acc. No. GCF_002902225). A marked divergence was observed between *S. sciuri* genomes enclosed within the MRSS C2865 sub-branch with the rest of *S. sciuri* genomes analyzed (96% ANI) (S2 Fig). A fined phylogenomic tree built using the core genome (1,331,382 bp) also evidenced the separation of these four strains into a separate branch within the species, approaching to the *S. vitulinus* and *S. fleuretti* cluster (Fig 1A). *S. sciuri* species enclosed isolates from human, animal, food, and environmental origin – including clinical isolates, while strains from the other group members were isolated from animal related sources.

**Figure 1.**
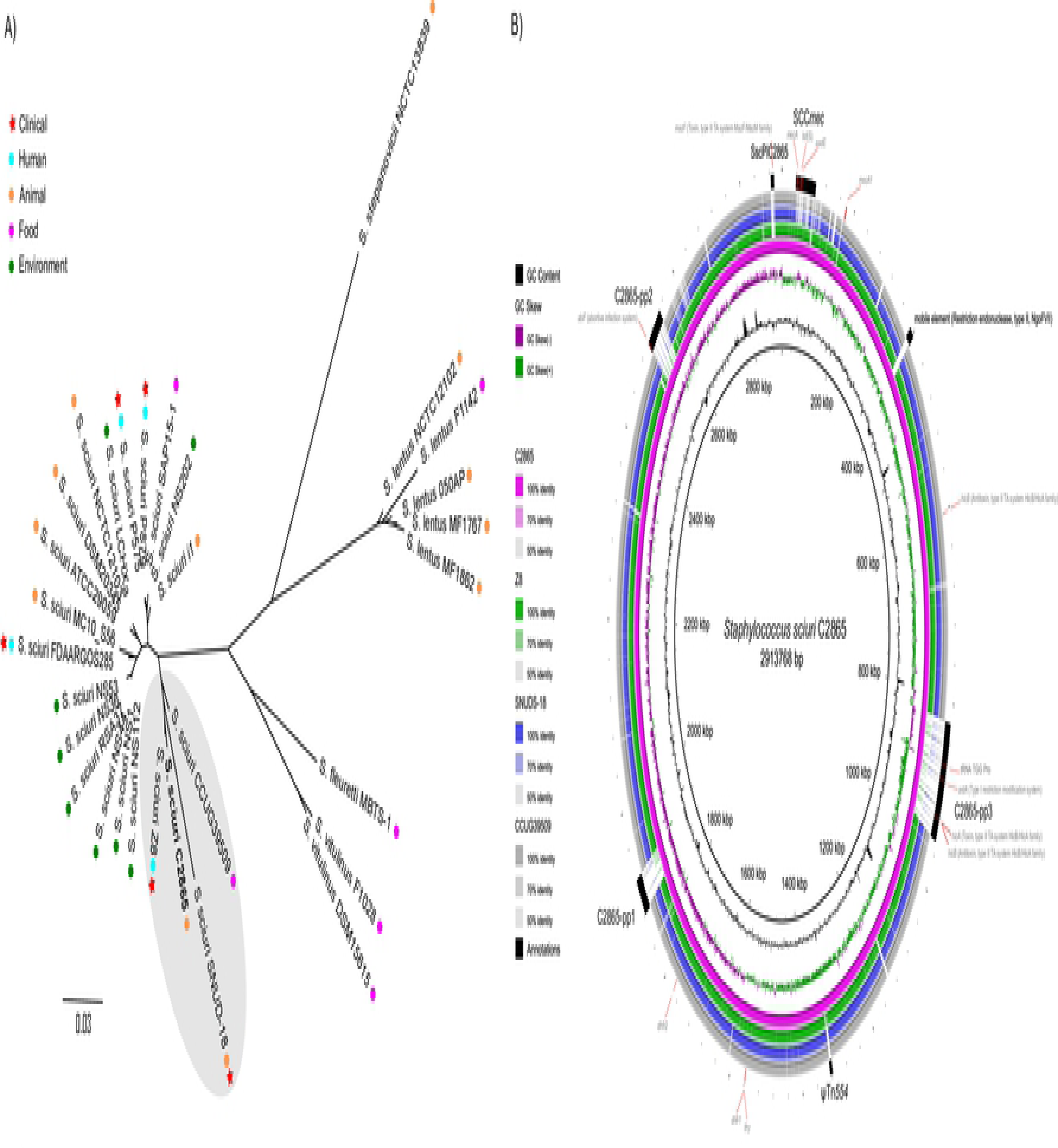
Phylogenomic analysis of *S. sciuri* group genomes and comparative diagram of MRSS C2865 closest relatives. A) Phylogenomic tree of *S. sciuri* C2865 and all 29 *Staphylococcus sciuri* group genomes deposited in the NCBI database (accessed until April 2018). Sample source is indicated as human, animal, food, environment as well as whether the sample was isolated from a clinical infection. Note: Strains *S. sciuri* DSM20345 and *S. sciuri* NCTC12103 actually correspond to the same type strain (also known as ATCC 29062). B) Comparative diagram of *S. sciuri* C2865 (pink ring) (reference genome) against its three closest genomes (*S. sciuri* Z8 [green ring], *S. sciuri* SNUD18 [blue], and *S. sciuri* CCUG39509 [grey]) as a set of concentric rings, where color indicates a BLAST match using Blast Ring Image Generator (BRIG) [77]. GC content and GC skew of reference genome is also displayed. Several features of interest are also depicted, highlighting the unique presence of remarkable mobile genetic elements (SCC *mec*, C2865-pp3, ψTn*554*, C2865-pp1, C2865-pp2, SscPIC2865) and several genes of interest in *S. sciuri* C2865.

A comparative diagram of MRSS C2865 chromosome against its three closest relatives (showing the regions unique to MRSS C2865) is depicted in Fig 1B. A number of chromosomal segments >5 kb unique to MRSS C2865 were detected, corresponding to novel recombinase site-specific MGEs (SCC*mec*, ψTn*554*, three prophages, staphylococcal pathogenicity island) described in the following sections. An extra region of ca. 11.5 kb was detected 390,146 bp downstream of *dnaA* (Fig 1B). This unique segment carried an NgoFVII family restriction endonuclease cluster (restriction modification system of the Type II), which was delimited at its downstream boundary by a transposase [N-terminal domain PF05598) and C-terminal domain DDE_Tnp_1_6 (PF13751) of the DDE superfamily]. This cluster included a Type II restriction endonuclease of the NgoFVII family (PF09562), which carries out the endonucleolytic cleavage of DNA to give specific double-stranded fragments with terminal 5’-phosphates (by recognizing the double-stranded sequence GCSG/C), and two consecutive DNA methyltransferases (DNA MTase) (PF00145).

Potential genes or chromosomal elements involved in conjugation, such as integrative or conjugative elements or conjugative transposons, which could mediate intercellular transfer of identified MGEs, were not detected. The genome location of the methicillin-susceptible *mecA1* gene, two intrinsic *dhfr* genes (*dhfr1* and *dhfr2,* the former next to a *thy* gene), a bacterial chromosomal *hicA*-antitoxin *hicB* (Type II TA system), as well as the most relevant adaptive genes from the novel chromosomally-located MGEs are displayed (Fig 1B).

### *S. sciuri* exhibits a highly open pan genome with MRSS C2865 showing the largest and most unique genome content

Whole-genome alignment of all *S. sciuri* genomes revealed a core genome of 1.3 Mb (58.9%) with 1,547 shared proteins at 95% of identity. The pan genome of 21 *S. sciuri* genomes (enclosing 5,721 genes) comprised the 1,547 genes (27%) forming the core genome of *S. sciuri* and 4,174 genes encompassing the accessory genome (73%) (Fig 2A). The pan genome curve revealed not saturated while the core genome was limited to less than 1,600 genes (Fig 2B). This indicates an open pan genome and high genetic diversity in *S. sciuri*. This is further supported by a high prevalence of “cloud genes” (genes found in up to 15% of the strains), which corresponds to approximately 43.2% of the pan genome (Fig 2A). Importantly, *S. sciuri* C2865 harbored the highest total gene number (3,063; average gene number 2,707) among the species, in addition to the highest number of unique genes (521; average gene number 95) (Fig 2C). Within this fraction, in addition to the chromosomally located elements mentioned above, three AMR resistance plasmids were detected. One of them, termed pUR2865-34, corresponded to a novel and unique mosaic MDR plasmid described in the section below. These chromosomal and extrachromosomal MGEs, including the ISs observed, represented 75% of C2865 unique genes (390/521), while 21% (110/521) could not be assigned any function. Significant differences were observed in the total number of pan-genome genes detected by MRSS C2865 PacBio and Illumina data (5,721/5,220, p-value=1.7E-06). These differences were associated with the accessory genome (4,174/3,584, p-value=2.1E-11) and unique genes (1,996/1,722, p-value=7.0E-06).

**Figure 2.**
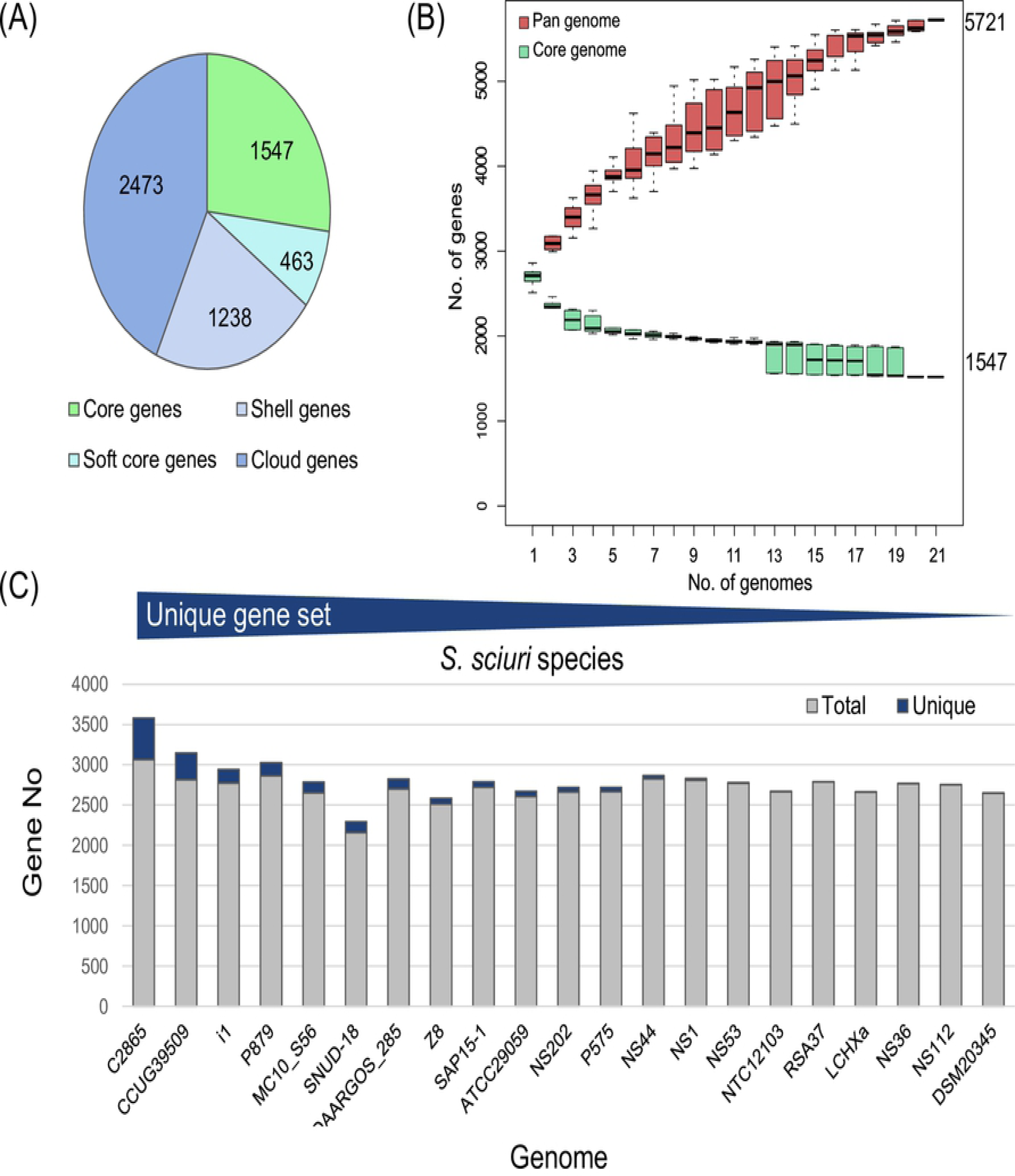
Pan genome analyses for *S. sciuri* genomes at species level (n=21, 95% identity). (A) Pie chart showing the proportions of coding DNA sequence (CDS) in the core, soft core, shell, and cloud genome. The parameters were defined as follows: Core genes: ≥ 99% of analyzed genomes, accessory genes: 1 - 99% (soft core 95-99%; shell 15-95%; cloud ≤ 15%). (B) Number of core genes (green) and total number of genes (pan genome) (blue) curve for 21 *S. sciuri* strains. The upper and lower edges of the boxes indicate the first quartile (25th percentile of the data) and third quartile (75th percentile), respectively, of 1000 random different input orders of the genomes. The central horizontal line indicates the sample median (50th percentile). (C) Bar chart of the total number of genes per genome indicating the amount of unique genes per genome (dark blue) ordered by the number of unique genes.

### Mosaic mobile adaptive elements enclosed within the novel plasmid pUR2865-34

Plasmid pUR2865-34 was 40,108 bp in size and harbored 41 open reading frames (ORFs) (Fig 3A). Illumina output plus scaffolding generated two potential plasmids (pUR2865-3 and pUR2865-4) flanked by IS*257* elements, which when combined together corresponded to pUR2865-34. Plasmid pUR2865-34 length and integrity was confirmed by plasmid linearization and gel electrophoresis in addition to WGS analysis of two *S. aureus* RN4220 transformants (RN4220/pUR2865-34) (data not shown). pUR2865-34 showed a mosaic IS*6*-like-delimited modular organization, consisting of novel and previously detected segments (Fig 3A), encompassing backbone and accessory regions. Four identical copies of staphylococcal IS*257* in the same orientation (with their 15-bp flanking perfect inverted repeats [IRs] 5’-GGTTCTGTTGCAAAG-3’) and one copy of the enterococcal IS*1216E* (with 23-bp flaking perfect terminal IRs 5’-GGTTCTGTTGCAAAGTTTTAAAT-3’) were identified.

**Figure 3.**
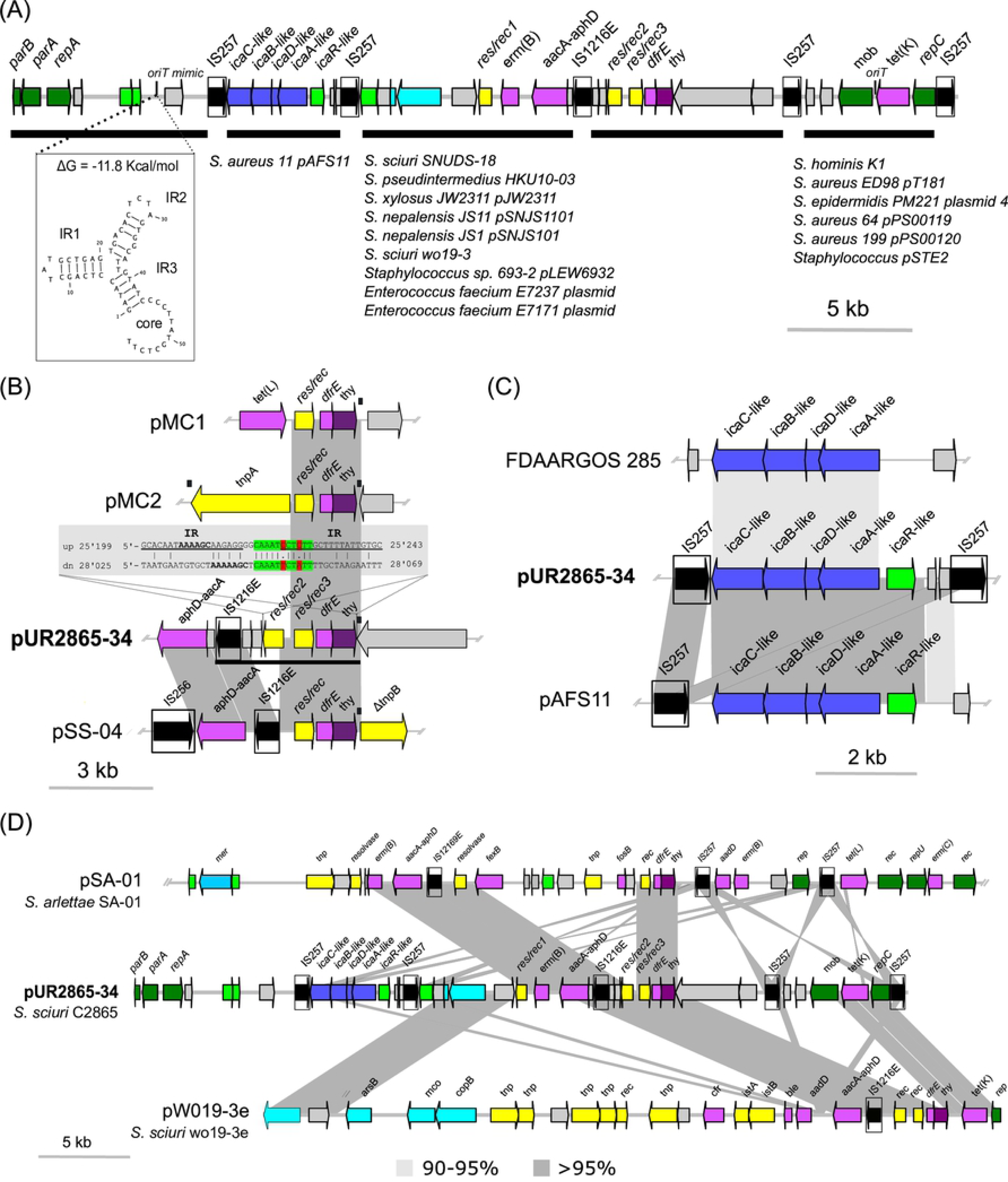
Comparative analysis of novel pUR2865-34 with the closest sequences in NCBI. Arrows denote the genes, length and orientation. Gene colors other than grey represent the following genes of interest (color): antimicrobial resistance genes (pink); genes involve in metal resistance or transport (bright blue); intercellular adhesion gene cluster (*ica*) genes (navy blue); genes involved in transcription regulation (bright green); plasmid backbone genes (dark green); genes involved in transposition or recombination (yellow) and insertion sequences with defined imperfect inverted repeats (IR) (boxed and black). Areas of nucleotide similarity (nblastn, >100 bp match, >80% ID) between strains/structures are indicated in grey. For plasmids pSA-01, pAFS11 and *S. sciuri* FDAARGOS plasmid unnamed only the area of interest is represented (A) *S. sciuri* C2865 pUR2865-34 underlining its modular organization and indicating strain/plasmid sequences detected with ≥50% coverage and ≥95% ID carrying those modules. The novel predicted secondary structure of the origin of conjugative transfer mimic (*oriT* mimic) is depicted, indicating the free energy of the DNA hairpins formation (ΔG) (B) Truncated *dfrE*-carrying transposon and immediate up- and downstream regions of *S. sciuri* C2865 pUR2865-34 and comparative area with *dfrE*-enclosing differing regions of *Exiguobacterium sp*. S3-2 pMC1 (GenBank acc. No. KF648874) and pMC2 (KF648875), as well as *S. sciuri* GN5-1 pSS-04 (KF129410). Potential up- and downstream sequence that might be involved in recombination is displayed and grey-shaded, with the precise plasmid position indicated. The 19-bp perfect IR (5’-GCACAATAAAAGCAAGAGG-3’) located upstream of *dfrE*-carrying remnant transposon are underlined. The 6-bp direct repeats (DR) (5’-AAAAGC-3’) and the 11-bp imperfect DRs (consensus: 5’-CAAAT[C/T]CT[C/A]TT-3’) located up- and downstream of *dfrE*-harboring region are bolded and highlighted in green with ambiguities in red, respectively. Black boxes above the graphical display represent the Tn*3*-like characteristic 38-bp inverted repeats detected, involved in excision and integration of Tn*3* related elements. Region conserved in all *dfrE*-carrying elements except for those graphically represented is underlined in pUR2865-34 segment. (C) *ica*-locus variant carrying region in *S. sciuri* C2865 pUR2865-34 and closest relatives deposited in NCBI database: *S. aureus* strain 11 pAFS11 (FN806789; displayed region 7’960 – 15’534 bp) and *S. sciuri* FDAARGOS plasmid (CP022047; 1 – 5’700 bp). (D) Represented plasmids closest to pUR2865-34 and strains carrying them are as follow (GenBank acc. no): *S. arlettae* strain SA-01 plasmid pSA-01 (KX274135; 1 - 42’500 bp) and *S. sciuri* strain wo19-3e plasmid (KX982172).

pUR2865-34 backbone region encompassed a replication initiation *repA* gene with the typical RepA_N N-terminal domain, characteristic of theta-replicating plasmids (data not shown) [43]. The N- and C-terminal domains of the encoded protein were homologous to different RepA_N proteins, reflecting the fact that these proteins can undergo domain replacement events. In addition, a Type Ib partitioning system, involved in plasmid segregational stability, consisting of a *parA* gene (coding for a deviant Walker ATPase motor protein) and a *parB* gene (coding for a DNA-binding protein with a RHH secondary structure motif), both located upstream of the *repA* gene, was detected. Resultant ParA and ParB shared highest identity with *Staphylococcus. aureus* apramycin resistance plasmid pAFS11 sequence, although the Type-Ib system was not annotated in this plasmid [44]. pUR2865-34 harbored a single relaxase gene (*mob*) with an origin of transfer (*oriT*) sequence immediately upstream this gene, which was enclosed within a rolling-circle replication (RCR) integrated plasmid, named pUR2865-int (Fig 3C). In addition, an *oriT* mimic sequence (positions 6,077 to 6,132) similar to one recently described by Bukowski *et al.,* [45], was observed proximal to the replication and partitioning genes (Fig 3A). However, as indicated above, conjugation related genes from known staphylococcal conjugative elements, such as pSK41, or integration and conjugative elements, which could act in *trans* for plasmid transference, were not detected throughout the entire bacterial genome.

Plasmid pUR2865-34 accessory region enclosed a copy of the tetracycline efflux MFS transporter gene *tet*(K), located within the small RCR integrated pUR2865-int. This plasmid belonged to the pT181 family, as identified by sequence homology of RepC (83.1%), the double strand origin (*dso*) region within *repC* (*dso nick* site: 5’-AAAACCGGaTACTCT/AATAGCCGGTT-3’) and the single strand origin (*ssoA*) region, located downstream of relaxase gene (Fig 3A). pUR2865-int shared 65.1% and 47.2% nucleotide identity to RCR prototype pT181 and pUR2865-2 (RepC 76.9% ID), respectively. This integrate was flanked by two identical copies of IS*257* in the same orientation. Two identical DRs on the internal ends (to the insertion) for each IS copy were detected, suggesting that this IS*257* set may have mediated insertion of a pT181-like ancestor carrying one IS copy with an IS*257*-carrying pUR2865-34 precursor by homologous recombination. Of note, the IS*257* copy adjacent to the *rep* gene in pUR2865-int has disrupted its promoter region (data not shown), rendering this gene likely not functional.

In addition, pUR2865-34 harbored a macrolide/lincosamide/streptogramin B resistance gene *erm*(B), coding for a 23S rRNA adenine N-6-methyltransferase and an aminoglycoside modifying *aacA-aphD* gene, coding for the bi-functional enzyme 6’-aminoglycoside N-acetyltransferase AAC(6’)-Ie aminoglycoside O-phosphotransferase APH(2’’)-Ia, located immediately downstream of an IS257/IS*1216E* copy in the same orientation.

Three S-rec genes of the resolvase/invertase subfamily, named *res/rec* (*res*/*rec1, res/rec2, res/rec3*), were identified. They contained the characteristic catalytic N-terminal PF00239 domain (with consensus motif 5’-Y-[AI]-R-V-<u>S</u>-[ST]-x (2)-Q-3’) and the C-terminal helix-turn-helix (HTH) domain of S-rec resolvases (PF02796). Res/Rec2 and /ResRec3 shared highest amino acid identity to limited staphylococcal isolates (four *S. aureus*, two *S. sciuri* and one *Staphylococcus arlettae* strains) (>93% identity), while most entries corresponded to resolvases found in other Gram positive bacteria (*Macrococcus caseolyticus, Exiguobacterium sp., Bacillus sp., Enterococcus sp., Lactococcus sp., Streptococcus sp., Lysinibacillus sp., Paenibacillus sp., Clostridium sp.*). A *dfr* gene, designated *dfrE*, and a thymidylate synthase gene (*thy*) were identified immediately downstream of *res*/*rec3*. Resultant DfrE, shared 31.5 and 15.5% identity to two additional intrinsic Dhfr detected in *S. sciuri* C2865 genome, both located in conserved chromosomal regions lacking ISs or other mobile elements. BLASTp searches revealed that DfrE was present in a limited number of strains at 100% sequence identity (Table 3). *dfrE*-carrying strains corresponded to all the staphylococcal isolates harboring the *ser*/*rec3* (see above) plus *M. caseolyticus* strain JCSC5402 and *Exiguobacterium* sp. strain S3-2, which enclosed the *dfrE* carrying region in two different coexisting plasmids (Table 3). All *dfrE*-enclosing staphylococcal genomes corresponded to WGS projects with direct submission to NCBI, without associated or linked publication. Remarkably, all *dfrE*-carrying genomes harbored it in MDR plasmids or plasmid-associated elements, hosting additional antibiotic and/or metal resistance genes (Table 3). Only Yang *et al.,* [46] denoted the *dfrE* gene in *Exiguobacterium* sp. S3-2 from a fish-farm sediment in China, which was resistant to TMP-sulfametoxazol (minimum inhibitory concentration [MIC] >1’024 µg/ml). The authors designated this gene as “*dfr_like*”, and experimentally proved its functionality in a Gram-negative heterologous host, *E. coli* DH5α, carrying the recombinant *pUC19/dfr_like* (MIC >2’048 μg/ml). Hence, the role of this gene - here *dfrE* (for *Exiguobacterium* spp) - in TMP resistance in staphylococci remained open.

**Table 3.**
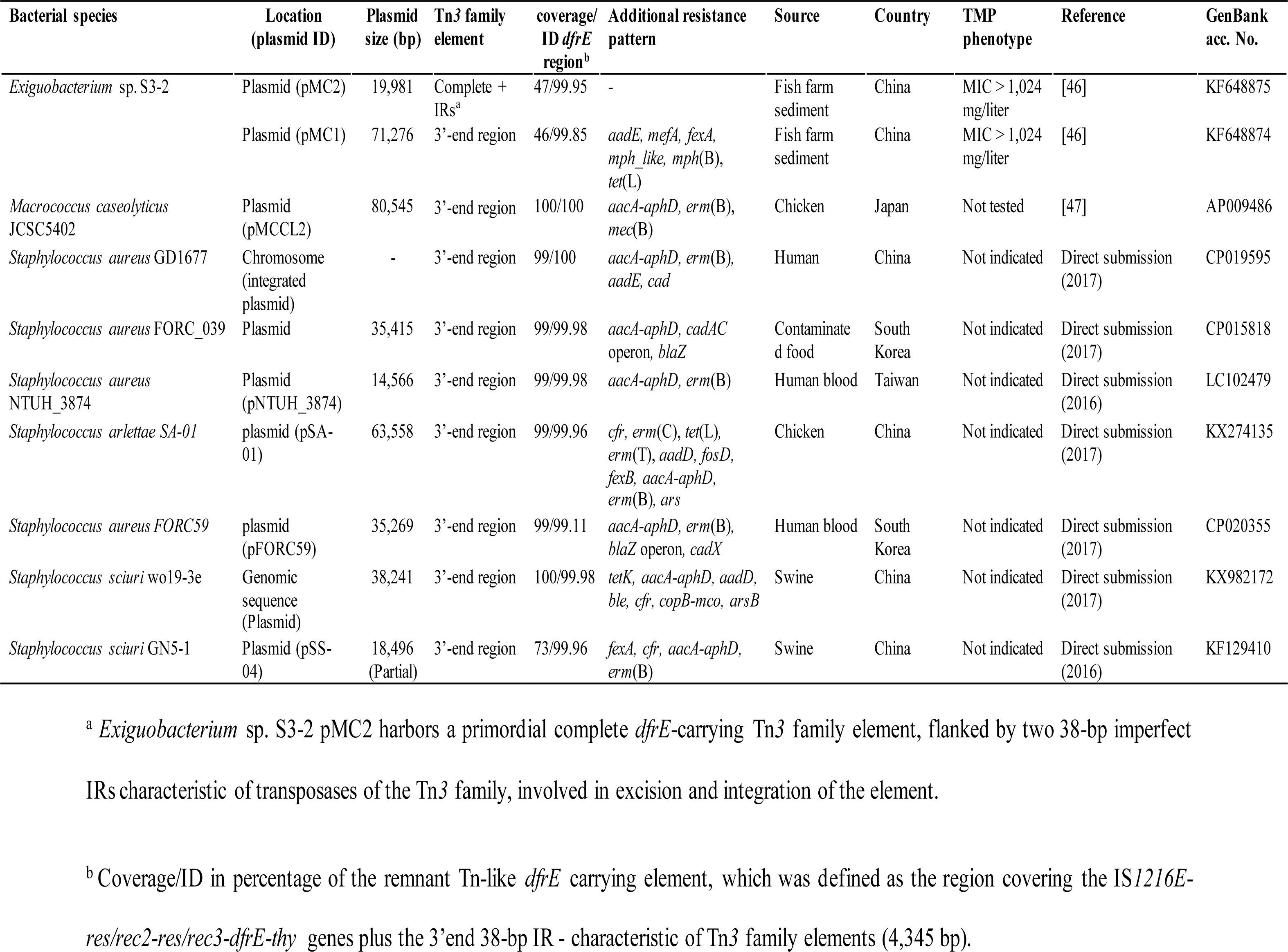
General features of *dfrE-carrying* strains deposited in the NCBI database and genetic platforms containing it.

| Bacterial species | Location (plasmid ID) | Plasmid size (bp) | Tn3 family element | coverage/ID *dfrE* region<sup>b</sup> | Additional resistance pattern | Source | Country | TMP phenotype | Reference | GenBank acc. No. |
| --- | --- | --- | --- | --- | --- | --- | --- | --- | --- | --- |
| *Exiguobacterium* sp. S3-2 | Plasmid (pMC2) | 19,981 | Complete + IRs<sup>a</sup> | 47/99.95 | - | Fish farm sediment | China | MIC > 1,024 mg/liter | [46] | KF648875 |
|  | Plasmid (pMC1) | 71,276 | 3'-end region | 46/99.85 | *aadE*, *mefA*, *fexA*, *mph_like*, *mph*(B), *tet*(L) | Fish farm sediment | China | MIC > 1,024 mg/liter | [46] | KF648874 |
| *Macrococcus caseolyticus* JCSC5402 | Plasmid (pMCCL2) | 80,545 | 3'-end region | 100/100 | *aacA-aphD*, *erm*(B), *mec*(B) | Chicken | Japan | Not tested | [47] | AP009486 |
| *Staphylococcus aureus* GD1677 | Chromosome (integrated plasmid) | - | 3'-end region | 99/100 | *aacA-aphD*, *erm*(B), *aadE*, *cad* | Human | China | Not indicated | Direct submission (2017) | CP019595 |
| *Staphylococcus aureus* FORC_039 | Plasmid | 35,415 | 3'-end region | 99/99.98 | *aacA-aphD*, *cadAC* operon, *blaZ* | Contaminated food | South Korea | Not indicated | Direct submission (2017) | CP015818 |
| *Staphylococcus aureus* NTUH_3874 | Plasmid (pNTUH_3874) | 14,566 | 3'-end region | 99/99.98 | *aacA-aphD*, *erm*(B) | Human blood | Taiwan | Not indicated | Direct submission (2016) | LC102479 |
| *Staphylococcus arlettae* SA-01 | plasmid (pSA-01) | 63,558 | 3'-end region | 99/99.96 | *cfr*, *erm*(C), *tet*(L), *erm*(T), *aadD*, *fosD*, *fexB*, *aacA-aphD*, *erm*(B), *ars* | Chicken | China | Not indicated | Direct submission (2017) | KX274135 |
| *Staphylococcus aureus* FORC59 | plasmid (pFORC59) | 35,269 | 3'-end region | 99/99.11 | *aacA-aphD*, *erm*(B), *blaZ* operon, *cadX* | Human blood | South Korea | Not indicated | Direct submission (2017) | CP020355 |
| *Staphylococcus sciuri* wo19-3e | Genomic sequence (Plasmid) | 38,241 | 3'-end region | 100/99.98 | *tetK*, *aacA-aphD*, *aadD*, *ble*, *cfr*, *copB-mco*, *arsB* | Swine | China | Not indicated | Direct submission (2017) | KX982172 |
| *Staphylococcus sciuri* GN5-1 | Plasmid (pSS-04) | 18,496 (Partial) | 3'-end region | 73/99.96 | *fexA*, *cfr*, *aacA-aphD*, *erm*(B) | Swine | China | Not indicated | Direct submission (2016) | KF129410 |

<sup>a</sup> *Exiguobacterium* sp. S3-2 pMC2 harbors a primordial complete *dfrE*-carrying Tn3 family element, flanked by two 38-bp imperfect
IRs characteristic of transposases of the Tn3 family, involved in excision and integration of the element.

Phylogenetic analysis of DfrE (>50% ID) in NCBI database plus all Dfrs responsible for TMP resistance in staphylococci revealed that DfrE was closer to the Dhfr of soil-related *Paenibacillus anaericanus* (68% ID) (Fig 4). Comparative analysis of (i) all Dhfr/Dfrs detected in the 30 *S. sciuri* group genomes available, (ii) all TMP resistance Dfrs formerly detected in staphylococci and (iii) two susceptible Dhfrs as reference (Dhfr from *S. epidermidis* ATCC12228 and DhfrB from *S. aureus* ATCC25923) revealed that both chromosomally located Dhfrs cluster with intrinsic, susceptible staphylococcal Dhfrs whereas DfrE shared closer phylogenetic identity with TMP resistance DfrF, typical of enterococci and streptococci (S3 Fig).

**Figure 4.**
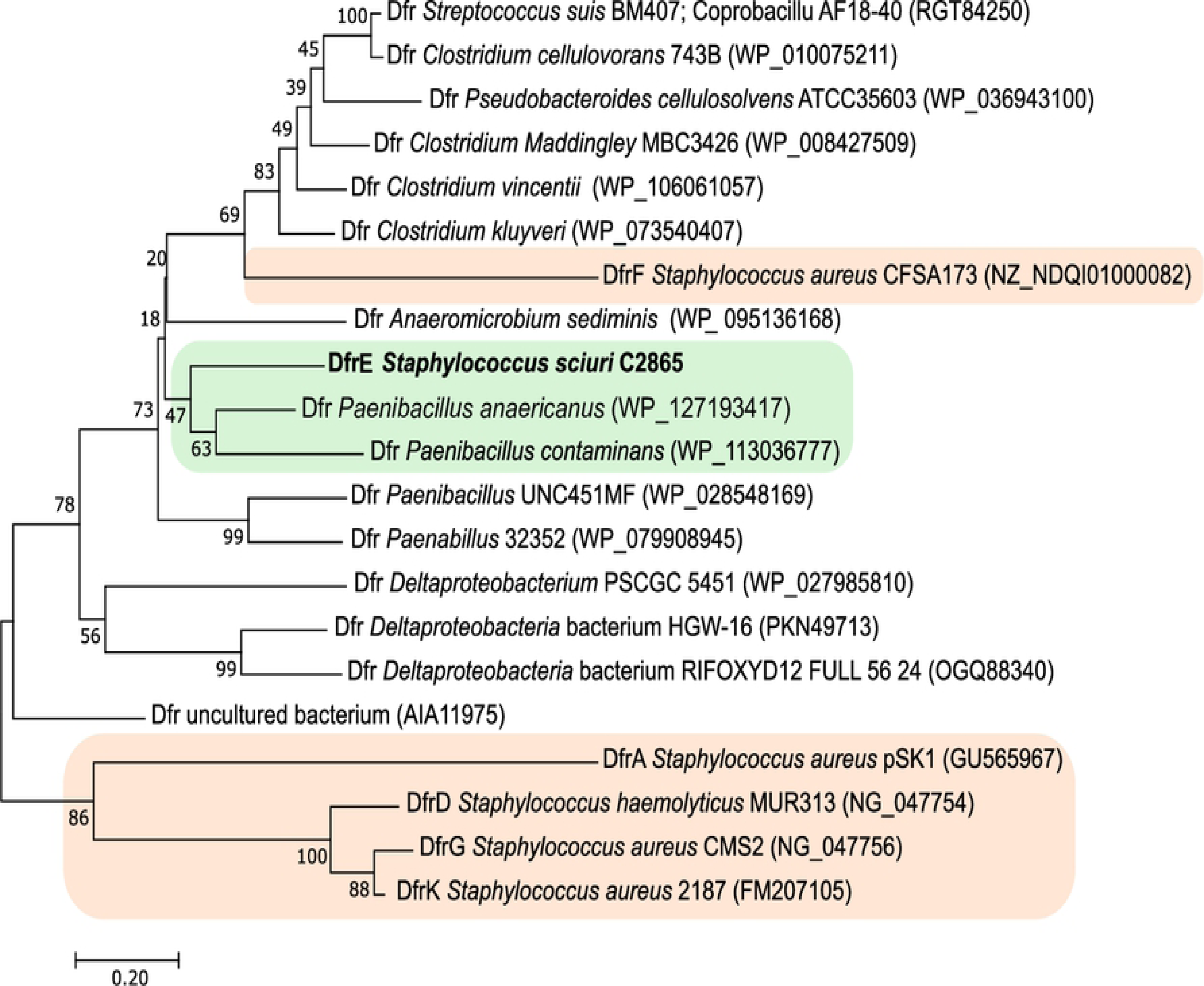
Neighbor-joining tree of aligned amino-acid sequences of the dihydrofolate reductases (Dfr) closest to the new DfrE. All Dfr proteins derived from BLASTp hits with a percentage of identity >50% to DfrE of *S. sciuri* strain C2865 were included, plus all trimethoprim resistance Dfrs described so far in staphylococci (DfrA, DfrD, DfrG, DfrK, DfrF) [35–39]. Identical Dfr proteins present in different bacterial classes were labelled with the ID of one representative per class (i.e. Firmicutes: *Streptococcus suis* BM407 and *Coprobacillus* sp. AF18-40). Gene, genera, species and strains (when available) are followed by their G enBank accession numbers within brackets. The trimethoprim resistant Dfr of *S. sciuri* C2865 is displayed in bold. The staphylococcal Dfr are backgrounded in faint orange, whereas the amino acid branch clustering the trimethoprim resistance DfrE is highlighted with a faint green background.

The *dfrE*-carrying region suggested a transposon-like structure encompassing an IS*1216E* copy, the *res/rec2* and *res/rec3* resolvase genes, as well as the *dfrE* and *thy* genes (4,345 bp) (Fig 3B). This element seems to be a truncated version of the original *dfr_like*-carrying transposon (Tnp) of the *Tn*3 family detected in plasmid pMC2 of *Exiguobacterium* sp. S3-2 [46], as only the 38-bp imperfect IRs downstream of *thy* was present (Fig 3B). Here, the pMC2-carrying Tn*3*-like *tnpA* gene, including the 38-bp upstream flanking IR, has been replaced by *res/rec2*. This truncated region (IS*1216E-res/rec2*-*res/rec3-dfrE-thy*) was conserved in all additional *dfrE*-enclosing strains (Table 3). Of note, two 6-bp identical DRs (5’-AAAAGC-3’) and two 11-bp imperfect DRs (consensus: 5’-CAAAT[C/T]CT[C/A]TT-3’), were detected immediately up- and downstream of this *dfrE*-carrying structure. In addition, two 19-bp perfect IRs (5’-GCACAATAAAAGCAAGAGG-3’) located upstream of *ser*/r*ec2* were detected, which might play a role in the recombination of this region (Fig 3B).

Plasmid pUR2865-34 harbored an additional IS*257*-flanked module, which encompassed a variant of the intercellular adhesion gene cluster (*ica*ADBC) (Fig 3C). These gene cluster consists of the intercellular adhesion operon (*ica*ADBC), comprising the *icaA, icaD, icaB* and *icaC* genes, and the adjacent *ica* locus repressor *icaR* gene, which is located upstream and transcribed in opposite direction from the *ica* operon [48]. The *ica* locus is involved in the early steps of biofilm formation (intercellular adhesion and cell agglutination) in *S. aureus*, *S. epidermidis* and another coagulase negative staphylococci. The *ica*-locus variant shared an overall 59.6% nucleotide identity with intercellular adhesion *ica*-locus prototype of methicillin-resistant *S. aureus* (MRSA) strain N315 (GenBank ac. No. BA000018) [44, 49]. Comparison of the deduced amino acid sequences with MRSA strain N315-encoded Ica proteins revealed identities of 60.9% for the IcaA protein, 35.8% for IcaD, 47.3% for IcaB, 47.2% for IcaD and as low as 26.3% for IcaR. This gene cluster shared highest identity (97.9%) to the *ica*-locus variant of *S. aureus* pAFS11, an apramycin resistance *apmA*-carrying plasmid (GenBank acc. No. FN806789.3) (Fig 3C), followed by the *ica*ADBC cluster present in *S. sciuri* FDAARGOS_285 chromosome and respective plasmid (CP022046.2 and CP022047.2, respectively). Regardless both IS copies likely mediated movement/capture of this adhesion cluster variant, IS*257*-characteristic 8-bp DRs were not detected.

MDR plasmids pSA-01 from *S. arlettae* strain SA-01 (KX274135) and pW019-3e from *S. sciuri* strain W019-3e (KX982172) resulted the closest relatives to mosaic pUR2865-34 (Fig 3D). These plasmids also contain the *dfrE*, *thy* and res/*rec3* carrying region.

### The novel *dfrE* gene alone confers high level resistance to trimethoprim

All four original *S. sciuri* strains exhibited high-level TMP resistance by MIC (≥4,096 µg/ml). *E. coli* DH5aα transformants harboring any of the three different *dfrE*-carrying constructs exhibited a 2,046 or 4,096-fold increase in TMP resistance with respect to the control (Table 4). Likewise, *S. aureus* RN4220 transformants enclosing the three different *dfrE*-carrying constructs exhibited a 2,048-fold increase in TMP resistance with respect to the empty *S. aureus* RN4220 and to reference *S. aureus* DSM 2569, irrespective of the presence of the natural or constitutive promoter located upstream *dfrE* (Table 4). Likewise, lack of synergistic activity was observed when *dfrE* and *thy* were cloned together into *E. coli* DH5α and *S. aureus* RN4220. Two *S. aureus* RN4220 transformants carrying entire pUR2865-34, named S319 and S320, exhibited 2,048-fold increase in TMP resistance with respect to reference and control strains, and displayed additional resistance to tetracycline, erythromycin, clindamycin, gentamicin, kanamycin, tobramycin, as confirmed by disc-diffusion agar tests.

**Table 4.** Minimum inhibitory concentration (MIC) values to trimethoprim (TMP) of original strains and respective DH5a and RN4220 constructs.

| Species | Strain | Characteristics, origin or description | Reference or source | Antimicrobial resistance gene/s | MIC<sup>c</sup><br>(μg/ml)<br>to TMP |
| --- | --- | --- | --- | --- | --- |
| *Escherichia coli* | DH5a | Recipient strain for electroporation, plasmid free | Promega | - | 0.5 |
|  | DH5a/pBUS1-HC | DH5a with *S. aureus-E. coli* shuttle vector pBUS1-HC | [50] | *tet*(L) | 0.5 |
|  | DH5a/pBUS1-Pcap-HC | DH5a with *S. aureus-E. coli* shuttle vector pBUS1-Pcap-HC | [50] | *tet*(L) | 0.5 |
|  | DH5a/1B-1 | DH5a / pBUS1-Pcap-HC / *dfrE* alone | This study | *tet*(L), *dfrE* | 2,048 |
|  | DH5a/2A-1 | DH5a / pBUS1-HC / + *dfrE* +<sup>a</sup> | This study | *tet*(L), *dfrE* | 2,048 |
|  | DH5a/4A-1 | DH5a / pBUS1-HC / + *dfrE* + *thy* | This study | *tet*(L), *dfrE* | 1,024 |
| *Staphylococcus aureus* | DSM 2569 | Reference strain for MIC (agar dilution method) | DSMZ<sup>b</sup> | - | 1 |
|  | RN4220 | Recipient strain for electroporation, plasmid free | DSMZ | - | 1 |
|  | RN4220/pBUS1-HC | RN4220 with *S. aureus-E. coli* shuttle vector pBUS1-HC | This study | *tet*(L) | 0.5 |
|  | RN4220/pBUS1-Pcap-HC | RN4220 with *S. aureus-E. coli* shuttle vector pBUS1-Pcap-HC | This study | *tet*(L) | 0.5 |
|  | RN4220/1B-1 | RN4220 / pBUS1-Pcap-HC / *dfrE* alone | This study | *tet*(L), *dfrE* | 2,048 |
|  | RN4220/2A-1 | RN4220 / pBUS1-HC / + *dfrE* + | This study | *tet*(L), *dfrE* | 2,048 |
|  | RN4220/4A-1 | RN4220 / pBUS1-HC / + *dfrE* + *thy* | This study | *tet*(L), *dfrE* | 2,048 |
|  | RN4220/pUR2865-34 | RN4220 with natural pUR2865-34 from *S. sciuri* C2865 | This study | *tet*(K), *erm*(B), *aacA-aphD*, *dfrE* | 2,048 |
| *Staphylococcus sciuri* | C2865 | Groin are of dog in Nigeria | [13] | *mecA*, *tet*(K), *tet*(M), *erm*(B), *lnu*(A), *aacA-aphD*, *ant6'*, *cat*<sub>pC221</sub>, *dfrE* | ≥ 4,096 |
|  | C2853 | Groin are of dog in Nigeria | [13] | *mecA*, *tet*(K), *erm*(B), *aacA-aphD*, *dfrE* | ≥ 4,096 |
|  | C2854 | Groin are of dog in Nigeria | [13] | *mecA*, *tet*(K), *tet*(M), *erm*(B), *aacA-aphD*, *cat*<sub>pC223</sub>, *dfrE* | ≥ 4,096 |
|  | C2855 | Groin are of dog in Nigeria | [13] | *mecA*, *tet*(K), *tet*(M), *erm*(B), *lnu*(A), *aacA-aphD*, *cat*<sub>pC221</sub>, *dfrE* | ≥ 4,096 |

<sup>a</sup> "+" represents *dfrE* immediate upstream or downstream region.
<sup>b</sup> Leibniz Institute DSMZ-German Collection of Microorganisms and Cell Cultures.
<sup>c</sup> MIC: Minimum inhibitory concentration.

### Ambiguous biofilm formation ability of *ica*-locus variant containing strains

Based on the CRAmod assay, all four *ica*-locus variant carrying *S. sciuri* strains were biofilm formers, while they were considered weak biofilm formers by the CV assay (S4 Fig). By this later method, both positive controls, strains SA113 and DSM1104, confirmed strong biofilm formers. Instead, remarkable lack of correlation was observed with the non-biofilm former *S. aureus* RN4220 and both RN4220 transformants carrying the pUR2865-34 plasmid (S319 and S320) between both assays (S4 Fig). Based on colorimetry by the CRAmod assay, neither recipient *S. aureus* RN4220 strain nor resultant transformants were biofilm formers (S4 Fig). Instead, by the CV assay, both recipient and resultant transformants resulted biofilm formers, with *S. aureus* RN4220 (S4 Fig). Yet, the *ica*-negative non biofilm producing *S. lentus* strain (C3030) confirmed lack of biofilm formation.

### *dfrE*, *ica*-locus variant and pUR2865-34-like are present in C2853, C2854 and C2855 strains

In addition to *S. sciuri* C2865, the three additional TMP resistant *S. sciuri* strains carried the novel *dfrE* gene and the *ica*-locus variant cluster, based on the PCR assays. Linearized-plasmid analysis by gel electrophoresis revealed that these three strains also harbored a similar-sized plasmid as pUR2865-34. Strains C2853, C2854 and C2855 were also resistant to tetracyclines, macrolides-lincosamides and aminoglycosides and harbored the corresponding *tet*(K), *erm*(B) and *aacA-aphD* resistance genes, which were also harboured in mosaic plasmid pUR2865-34 (Table 4).

### Small single resistance rolling-circle replicating pUR2865-1 and pUR2865-2 plasmids

Small plasmids pUR2865-1 (2,559 bp) and pUR2865-2 (3,830 bp), which carried the *lnuA* gene, encoding the lincosamide nucleotidyltransferase LnuA, and the *cat*_pC221_ gene, encoding the chloramphenicol acetyltransferase Cat_pC221_, respectively, were detected (Fig 5A, S5 Fig). Both plasmids consist of a basic plasmid backbone, with a replication initiation protein (Rep), as well as a mobilization initiation protein or relaxase (Mob) in the case of pUR2865-2. Here, an *oriT* was detected in the immediate upstream region of *mob*. Both *dso* and *sso* were detected in both plasmids based on sequence homology and secondary structure analysis (Fig 5B-C).

**Figure 5.**
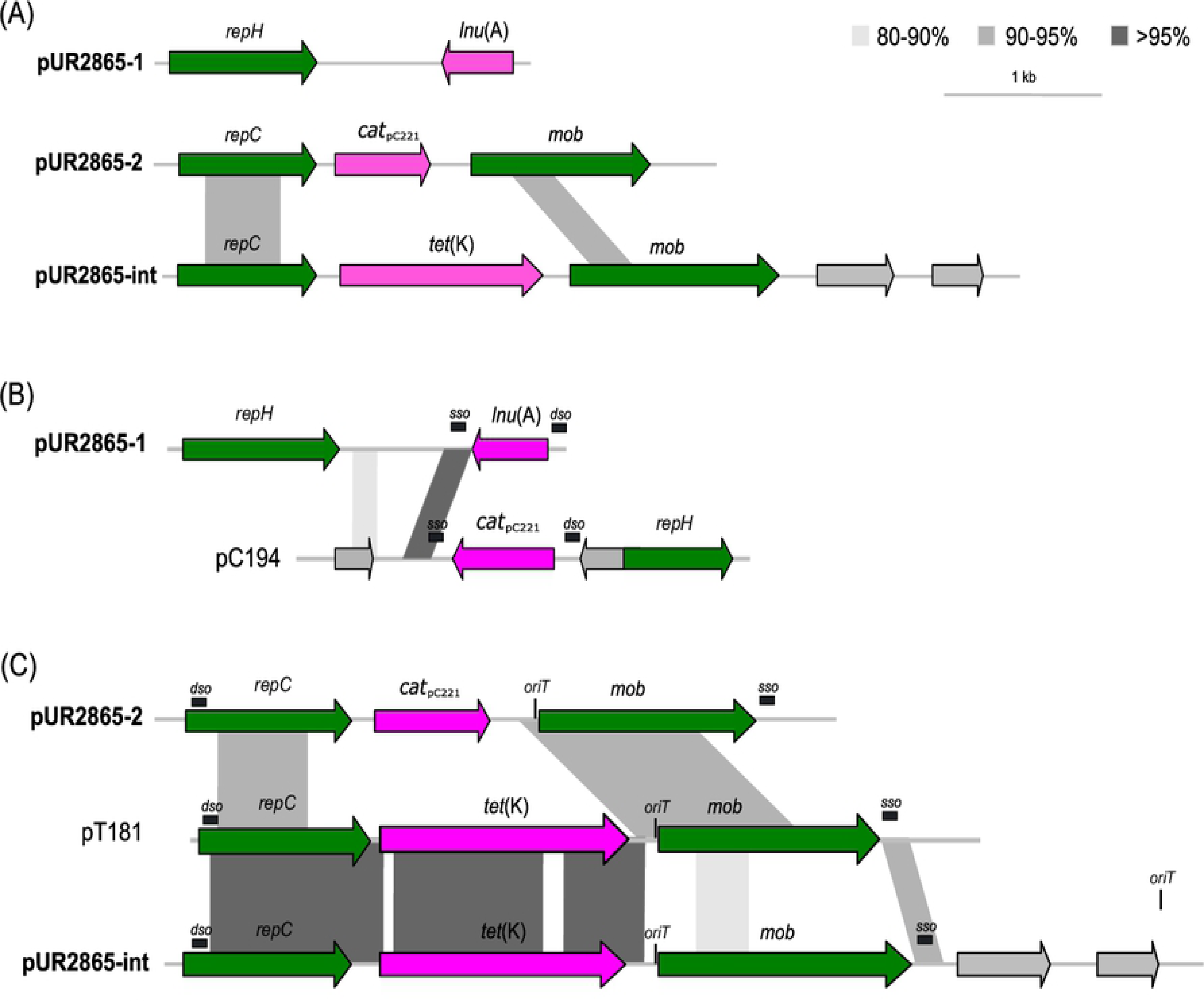
Graphical representation of the RCR plasmids detected in MRSS C2865 and comparative analysis with representatives of RCR family members. (A) Nucleotide sequence comparative analysis of the three 3 RCR plasmid s detected in MRSS C2865: pUR2865-1, carrying the lincosamide resistance *lnu*(A) gene, pUR2865-2 harboring the chloramphenicol resistance gene *cat*pC221, and pUR2865-int, which is integrated in the larger pUR2865-34 plasmid and carries the tetracycline resistance gene *tet*(K). (B) Nucleotide sequence comparative analysis of pC194-related pUR2865-1 and *S. aureus* pC194 (Genbank acc. No. NC_002013). Estimated double strand origin (*dso*) and single strand origin (*sso*) of replication, based on sequence homology with those of pC194, are indicated. (C) Nucleotide sequence comparative analysis of the pT181-related pUR2865-2, the integrated pUR2865-int and the RCR family prototype *S. aureus* pT181 (Genbank acc. No. J01764.1). Putative *dso nick* site for pUR2865-1 and pUR2865-2 were 5’-TCTTCTTgTCTTG/A TAcTA-3’ (capital letters denote conserved bases with respect to pC194, and slash the site of cleavage) and 5’-AAAACCGGCgTACTCT/AATAGCCGGTT-3’, respectively, and positions are denoted [106]. Location of the characteristic *ssoA*-enclosed conserved recombination site (RSB) 5’-GAGAAAA-3’ and the primer RNA transcriptional terminator CS-6 (5’-TAGCGT-3’), needed for replication initiation of the lagging strand are also indicated. Origin-of-transfer (*oriT*) for pT181-related plasmids is also indicated.

Based on the *dso* region, amino acid identity and motif analysis of Rep with one prototype plasmid per RCR family [43], the novel plasmid pUR2865-1 belonged to the pC194 family (Fig 5B). Even though Rep identity to prototype pC194 was substantially low (Rep 24.1% ID), all pC194-family conserved motifs were present in pUR2865-1, highlighting the HUH motif (whose His residues are involved in metal ion coordination required for the activity of RCR) and the HUH C-proximal Glu213 and Tyr217, corresponding to both catalytic residues [51]. Plasmid pUR2865-2 shared clear conserved regions with the pT181 family (Rep 69.4% ID) (Fig 5C). For both plasmids, the putative *dso nick* site and *ssoA*-enclosed conserved regions, involved in replication of the leading and lagging strand, respectively, were detected and are denoted in Fig 5.

Comparative analysis of pUR2865-1 with closest publicly available plasmids is shown in S5A Fig. Such plasmids were present in different staphylococcal species (*S. aureus*, *Staphylococcus chromogenes*, *Staphylcococus simulans*, etc) of diverse origins (human and dairy cattle clinical samples). As for pUR2865-2, closest relatives shared 70-78% coverage and ≥ 81% nucleotide ID, and they were mainly enclosed in *S. aureus* of different origins (S5B Fig).

### Novel SCC*mec*_C2865_ element lacking formerly described chromosomal cassette recombinases

Two chromosomally located homologues of the *mecA* gene were detected : the methicillin - resistance determinant *mecA* and an intrinsic copy of the methicillin-susceptible variant *mecA1*. Resultant proteins shared 81.4% identity. The *mecA1* gene of *S. sciuri* C2865 shared 99.7% nucleotide identity (99.4% at amino acid level) to the *mecA1* gene present in methicillin - susceptible *S. sciuri* strain K11 (GenBank acc. No. Y13094), exhibiting five-point substitutions. One of these substitutions implied a stop codon (Glu607Stop) at the 3’-end region of penicillin binding protein transpeptidase domain (PF00905), resulting in a truncated shorter variant of MecA1. Both, *mecA* and *mecA1*, were located on the chromosomal right-arm near the start origin of replication (*dnaA*) (41,464 and 177,170 bp downstream of *dnaA*, respectively).

The *mecA* gene, conferring methicillin resistance, was located at the 3’-end region of 23S rRNA (pseudouridine(1915)-N(3))-methyltransferase RlmH gene (*rmlH*), which encloses the integration site of SCC*mec* elements. A novel MDR SCC*mec* cassette, denominated SCC*mec*C2865, was detected at this site. SCC*mec*C2865 was 55,137 bp in size and contained a class A *mec* gene complex (IS*431-mecA-mecR1-mecI*) (Fig 6A). Importantly, none of the so far described staphylococcal or macrococcal chromososome cassettes recombinase genes (*ccr*), which resultant LSR proteins are responsible for excision and integration of the cassette, were detected. SCC*mec*C2865 was delimited at both ends by characteristic SCC*mec*-flanking DRs with typical insertion site sequences (ISS) or attachment sites (*att*): *att*R 5’-GAAGCATATCATAAATGA-3’ at the 3’-end of *rlmH* and an identical direct repeat (DR) designated *att*L3 at the right boundary of the cassette, defining a transferable unit. Two additional imperfect *att* sites (*att*L1 5’-GAAGCGTATCACAAATAA-3’ and *att*L2 5’-GAGCCATATAATAAATAA-3’) were detected within SCC*mec*C2865 at base-pair position 29,884 and 41,929 of the cassette, respectively. This ISS distribution configures six units with the potential to form circularly excised elements: three different SCC*mec* cassettes (segments enclosing the *att*R + either *att*L1, *att*L2 or *att*L3) and three SCC cassettes (*att*L1 + either *att*L2 or *att*L3; *att*L2 + *att*L3). (Fig 6A). Characteristic imperfect IRs, required for CcrAB or CcrC recognition of the *att* sites, were located at the internal boundaries of the cassette (Fig 6A) [52].

**Figure 6.**
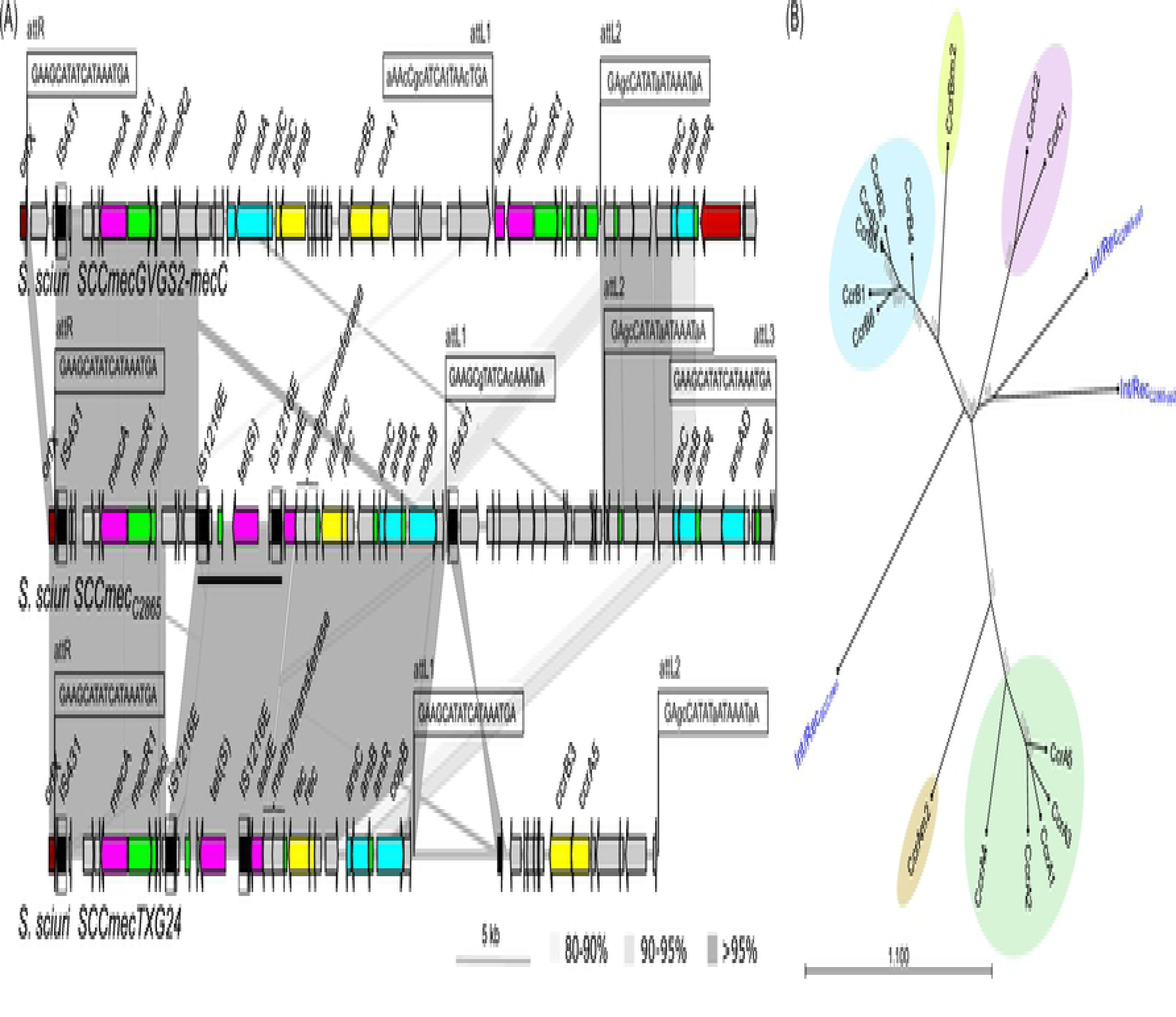
Graphical comparative analysis of the novel SCC *mec* and related large serine recombinases (LSR) detected in MRSS C2865. (A) Novel SCC*mec*C2865 (55,137 bp) and its closest elements in NCBI [SCC*mec*GVGS2-*mecC* of *S. sciuri* strain GVGS2 (GenBank acc. No. HG515014) and SCC*mec*TXG24 of *S. sciuri* strain TXG24 (KX774481) [15, 23]. Arrows denote the genes, length and orientation. Gene colours other than grey represent the following genes of interest (colour): antimicrobial resistance genes (pink); genes involve in metal resistance or transport (bright blue); genes involved in transcription regulation (bright green); genes involved in transposition or recombination (yellow); insertion sequences with defined imperfect inverted repeats (boxed and black); phage related genes (red) and the SCC *mec* integration gene (dark red), which strictly does not belong to the SCC*mec* cassette, except for the *rlmH* 3’-end terminal 18-bp attachment site (ISS1 or *att*R 5’-GAAGCATATCATAAATGA-3’). The two perfect direct repeats found at both extremities of the cassette (*att*R 5’-GAAGCATATCATAAATGA-3’ and *att*L3 5’-GAAGCATATCATAAATGA-3) are depicted. Two additional imperfect *att* sites (*att*L1 5’-GAAGCGTATCACAAATAA-3’ and *att*L2 5’-GAGCCATATAATAAATAA-3’) within SCC*mec*C2865 at base-pair position 29,884 and 41,929 of the cassette, respectively, are also depicted. Attachment sites detected in SCC*mec*GVGS2-*mecC* and SCC*mec*TXG24 are indicated. Unique bases with respect to *att*R, per SCC*mec* cassette, are represented in lower case. Areas of nucleotide similarity (nblastn, >100 bp match, >80% ID) between SCC*mec* cassettes are indicated in grey scale. (B) Neighbor-joining tree of aligned amino acid sequences of one representative staphylococcal chromosomal cassette recombinase (Ccr) per allotype described in *Staphylococcus* spp. and in *Macrococcus* spp. as well as the three LSRs present in *S. sciuri* C2865 genome (colored in blue) comprising the Ccr consensus motif Y-[LIVAC]-R-[VA]-S-[ST]-x(2)-Q or Y-[LIVAC]-R-[VA]-S-[ST]-x(4)-Q, the latter for the LSR detected within SCC*mec*C2865. LSR detected in *S. sciuri* C2865 encompassing the abovementioned motifs originate from SCC*mec*C2865 (Int/RecSCC*mec*); prophage vB_SscS_C2865-pp1 (Int/RecC2865-pp1) and prophage vB_SscS_C2865-pp2 (Int/RecC2865-pp2).

The SCC*mec*C2865 *rlmH*-proximal region, flanked by *att*R and *att*L1, stretched a region of ca. 31 kb that carried the class A *mec* gene complex, and one transposon-like structure consisting of one tetracycline resistance gene *tet*(S), coding for a translation elongation factor G (EF-G) involved in tetracycline ribosomal protection, flanked by two IS*1216E* copies in the same orientation. No DRs were present evidencing its mobilization as a composite Tn or translocatable unit. The internal IS*1216E*-flanked *tet*(S)-carrying segment was common among different Gram-positive bacteria, and shared highest nucleotide identity with corresponding regions in *Carnobacterium divergens* plasmid pMFPA43A1405B (LT984412), *Macrococcus sp*. IME1552 (CP017156), and *Lactococcus lactis* UC08 and UC11 plasmids pUC08B and pUC11B, respectively (CP016727 and CP016721), in addition to *S. sciuri* TXG24 SCC*mec*TXG24 (Fig 6A). A streptomycin resistance gene *aadE*, coding for a 6-aminoglycoside adenylyltransferase, was detected immediately downstream of the 3’-end IS*1216E* copy. Two consecutive genes in the same orientation coding for S-rec were identified. Protein domain analysis revealed that designated Int/RecSCC*mec* was a LSR, encompassing the phage-related typical PF00239, PF07508 and PF13408 domains, also present in Ccrs. In addition, an arsenic resistance operon (*arsCBR*) and a copper resistance gene (*copB*), coding for a copper-translocating P-type ATPase, were located within the *mecA*-carrying transferable unit (*att*R-*att*L1) (Fig 6A). SCC unit within *att*L1-*att*L2 boundaries consisted of a unique 11-kb region enclosing a cluster of genes coding for several metabolic routes. Of note, part of this region (62-66% coverage, >96% ID) was present in the chromosome of *S. xylosus* SA4009, *S. xylosus* S04010 and *S. xylosus* C2a [53], just after the SCC*mec* downstream junction. The *att*L2-*att*L3 module encompassed another copy of the arsenic resistance operon CBR (*arsCBR*) and an additional arsenic resistance gene (*arsAD*) plus an arsenic transcriptional regulator. SCC*mec*TXG24 from *S. sciuri* TXG24 and SCC*mec*GVGS2 from *S. sciuri* GVGS2 were identified as closest relatives to SCC*mec*C2865, sharing 57% and 40% coverage, respectively [15, 23]. Both cassettes harbored different allotypes of the typical SCC*mec ccrA* and *ccrB* genes.

It has been shown that Ccrs can still transpose SCC*mec* elements when located elsewhere in the genome [54]. Hence, the possible presence of *ccr* gene variants outside SCC*mec*C2865 was determined by a search of the translated genome of MRSS C2865 for the site-specific Ccr S-rec motif Y-[LIVAC]-R-[VA]-S-[ST]-x(2)-Q) present in LSR. Two hits were identified: (i) Int/Rec from prophage vB_SscS-C2865-pp1, designated Int/Rec_C2865-pp1_ and (ii) Int/Rec from prophage vB_SscS-C2865-pp2, designated Int/Rec_C2865-pp2_ (see below). Int/Rec of SCC*mec*C2865 (Int/RecSCC*mec*) was identified only when searching for consensus motif Y-[LIVAC]-R-[VA]-S-[ST]-x(<u>4</u>)-Q). Phylogenetic analysis of one representative Ccr per staphylococcal and macrococcal SCC*mec*, plus these three LSRs disclosed that they remarkably differ from staphylococcal CcrA, CcrB and CcrC allotypes and from macrococcal CcrAm2 an CcrBm2. Yet, Int/Rec_C2865-pp1_ and Int/Rec_C2865-pp2_ revealed phylogenetically closer to CcrCs (Fig 6B). Closest LSR to currently described Ccrs was Int/Rec_C2865-pp1_ (sharing 27.4% ID with CcrC2 and CcrB4), followed by Int/Rec_C2865-pp2_ (25.2% ID with CcrB4) and Int/RecSCC*mec* (24% identity with CcrC1). Of note, PCRs for potential CIs of the different units, as well as an intact chromosomal *rlmH* region (after SCC*mec* excision) were negative.

### Chromosomal *radC* integration of cadmium resistance ψTn*554*

A variant of the MLSB and spectinomycin resistance unit transposon Tn*554*, known as ψTn*554*, was identified integrated in the chromosome truncating the DNA repair *radC* gene at its 3’ region, 1’316’939 bp downstream of *dnaA* gene (S6 Fig). This ψTn*554* was 7’306 bp and belonged to the Tn*554* family of unit transposons, which insert at high efficiency into a primary unique site in the staphylococcal chromosomes (*att554*, within the *radC* gene) [43]. Transposon ψTn*554* contains a cadmium resistance gene cluster whose products code for a predicted transcriptional regulator (CadC), a cadmium-translocating P-type ATPase (CadA) and a cadmium resistance transporter family protein (CadD). ψTn*554* shared highest identity (94% coverage, 97.5% identity) with a *radC*-independent chromosomal region detected in a highly cytotoxic and clinically virulent *S. aureus* strain 6850 [55] (S6 Fig). Within *S. sciuri* available genomes, only a truncated version of this ψTn*554* was present within SCC*mec*GVGS2 cassette of *S. sciuri* strain GVGS2 (Fig 6), representing the single former report of a ψTn*554*-like structure in this species [23]. Nine additional *S. sciuri* genomes revealed to carry interrupted *radC* copies or 3’-end *radC* remnants (data not shown). In all these cases, the *radC* gene was intruded by a mobile element carrying the recently described oxazolidinone and phenicol resistance gene *optrA* (S6 Fig).

### Unique *Siphoviridae* prophages vB_SsS-C2865-pp1, vB_SsS-C2865-pp2 and vB_SsS-C2865-pp3 enclose adaptive features and excision and circularization ability

Three novel prophages were identified integrated at different positions of the chromosomal DNA of MRSS C2865. According to viral naming guidelines recommended by Adriaenssens *et al.,* [56], and following the naming system proposed by Kropinski *et al.,* [57], these new prophages were named as follows: vB_SsS-C2865-pp1 (41,284 bp), vB_SsS-C2865-pp2 (45,020 bp) and vB_SsS-C2865-pp3 (126,192 bp). Figs 5 and 6 show a comparative analysis of prophages C2865-pp1 and C2865-pp2, and C2865-pp3, respectively, with their closest relatives, In addition, phylogenetic analysis of the closest integrases (>70% coverage, >80% ID), all recovered from putative (pro)phages is displayed. The integrases from C2865-pp1 and C2865-pp2 encompassed the characteristic LSR resolvase, recombinase and the zinc beta ribbon domains (PF00239, PF07508 and PD13408, respectively), and, as indicated above, shared the core consensus motif typical of LSR Ccrs from SCC*mec* elements (Y-[LIVAC]-R-[VA]-S-[ST]-x(2)-Q) (Fig 6B). Sequencing analyses of potential CIs as well as restorage of respective phage chromosomal integration genes revealed that the three prophages had the ability to excise the bacterial genome and a single copy of the initial flanking DR was present in the resultant CI (*attP*) while the other remained in the restored phage integration gene (*attB*) in the bacterial genome.

#### C2865-pp1

C2865-pp1 was 41,283 bp long and consisted of 62 coding sequences (CDSs) of which 26 (41.9%) had predicted functions. C2865-pp1 had a GC content of 34.6% and was integrated truncating a VOC family-protein gene, 1,880,677 bp downstream of *dnaA*. The structural gene distribution of C2865-pp1 shared typical phage modular organization and characteristic phage gene groups involved in lysogeny (integrase, represors, antirepresor), DNA metabolism (single strand DNA binding, methyltransferase), packaging (HNH endonuclease, terminases: TerS, TerL), morphogenesis (major capsid, tail fiber, tape measure), and cell lysis (peptidase, holin, endolysin) were detected from left to right arm (Fig 7). The entire genome, with the exception of three genes of the lysogenic region, was rightwards transcribed.

**Figure 7.**
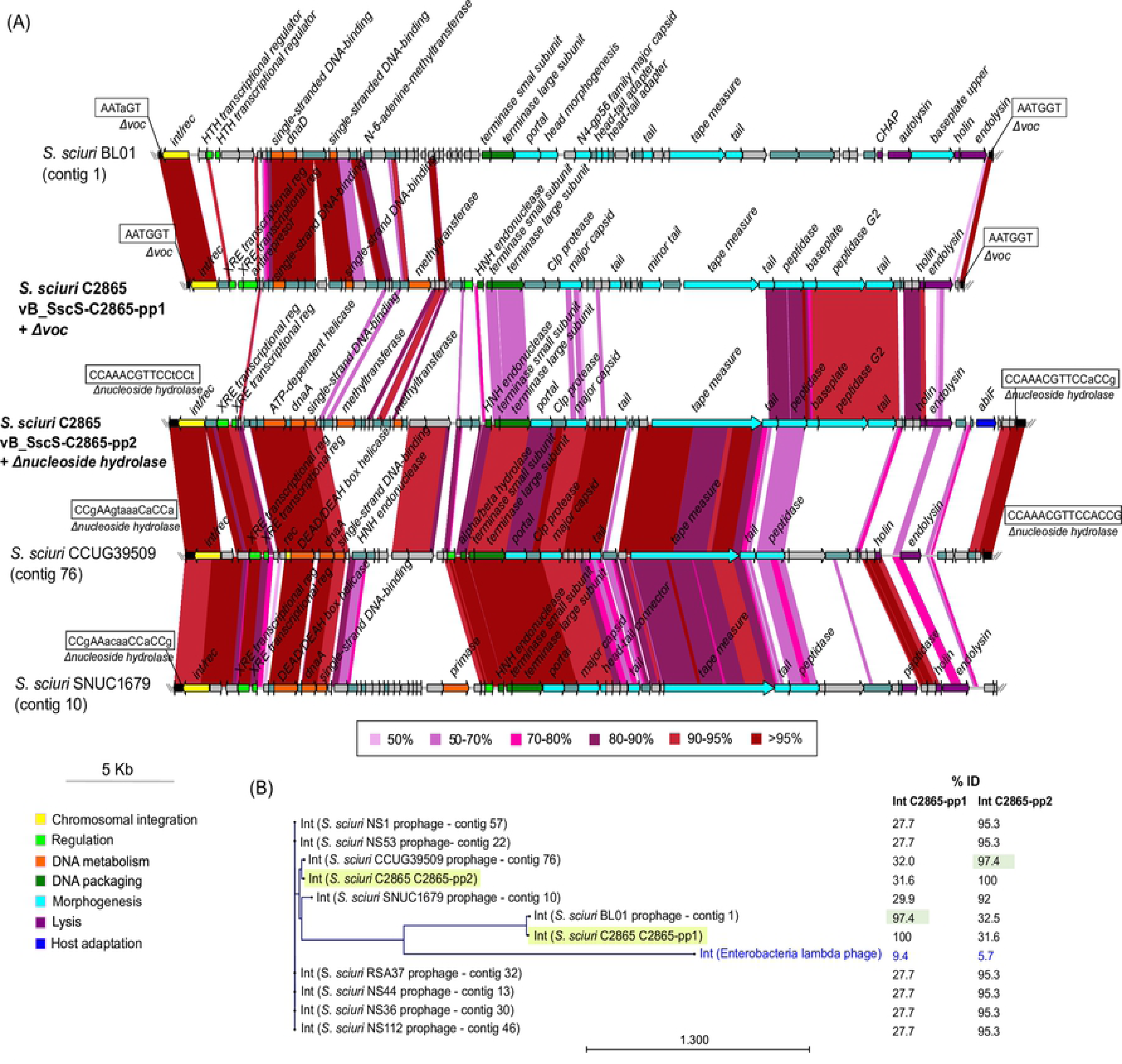
Graphical comparative analysis of prophages C2865-pp1 and C2865-pp2 and closest relatives. (A) Graphical comparative analysis of the novel staphylococcal prophages vB_SscS-C2865-pp1 (C2865-pp1) and vB_SscS-C2865-pp2 (C2865-pp2) integrated in the chromosome of *S. sciuri* strain C2865 and its closest relatives based on (i) genetic elements enclosing the closest integrases in the NCBI pBLAST database (*S. sciuri* BL01– contig 1 – [WGS, RefSeq acc. No. NZ_SPPH01000001], *S. sciuri* CCUG39509 – contig 76 – [NZ_PPRP01000076], *S. sciuri* SNUC1679 – contig 10 – [NZ_QXVB01000010]), as well as on (ii) all staphylococcal phages deposited in the Viral RefSeq database (accessed until November 2018). Arrows denote the genes, length and orientation. Gene colors other than grey (hypothetical proteins) and grey-turquoise (others) represent genes involved in the following processes (color): chromosomal integration (yellow), regulation (green), DNA metabolism (orange), DNA packaging (dark green), phage morphogenesis (bright blue), cell lysis (purple), host adaptation (navy blue). Areas of identity (tblastx, >50 aa match, >50% ID) between (pro)phages are indicated in color scale. (B) Maximum likelihood phylogenetic tree of C2865-pp1 and C2865-pp2 integrases, as well as all integrases sharing >80% amino acid identity (pBLAST search) with respect to either C2865-pp1 and C2865-pp2 integrases (*S. sciuri* NS1 - contig 57 - [WGS, NCBI acc. No. LDTK01000057], *S. sciuri* NS53 - contig 22 - [LDTP01000022], *S. sciuri* CCUG39509 - contig 76 - [WGS, RefSeq acc. No. NZ_PPRP01000076], *S. sciuri* SNUC1679 - contig 10 - [NZ_QXVB01000010], *S. sciuri* BL01 - contig 1 - [NZ_SPPH01000001], *S. sciuri* RSA37 - contig 32 - [LDQ01000032], *S. sciuri* NS44 - contig 13 - [LDTO01000013], *S. sciuri* NS36 - contig 30 - [LDTN01000030], *S. sciuri* NS112 - contig 46 - [LDTL01000046]), and integrase of Enterobacteria lambda phage (NC_001416), used as outgroup (colored in blue). C2865-pp1 C2865-pp2 integrases are highlighted in faint yellow. Percentage of identity of all integrases analyzed is indicated, with that sharing highest identity to respective integrase in faint green background.

The LSR integrase gene, named *int*/*rec*C2865-pp1, was detected at the left extremity of the phage. Two transcriptional regulators of the XRE superfamily (PF01381) were detected in front of *int*/*rec*_C2865-pp1_. These genes are homologs of lambda phage transcriptional regulators *cI* and *cro* (Cro/C1-type HTH domain) responsible for maintenance of phage integration (CI, lysogeny) and excision (Cro, lytic cycle). In addition, a phage antirepresor was identified immediately downstream of the second transcriptional regulator. Chromosomal integration of C2865-pp1 generated a 6-bp perfect DR (*attL* and *attR*) (5’-AATGGT-3’) (Fig 7) at the boundaries of C2865-pp1. Comparative analysis of Int/RecC2865-pp1 on the NCBI protein database revealed a single hit enclosed in *S. sciuri* strain BL01 (NCBI RefSeq. No. NZ_SPPH01000001). This integrase belonged to a putative prophage integrated at the same chromosomal gene as C2865-pp1; however, the DR were not exactly conserved, with consensus sequence 5’-AAT[A/G]GT-3’(Fig 7B).

#### C2865-pp2

C2865-pp2 was 45,020 bp and consisted of 64 CDSs of which 27 (42.2%) had predicted functions. C2865-pp2 had GC content of 33.8% and was integrated truncating a nucleoside hydroxylase gene, 2,501,758 bp downstream of *dnaA*. C2865-pp2 CDSs were arranged in functional modules in synteny with those detected in C2865-pp1. Most of the genome, except for the left-arm lysogenic region (3 CDSs) and the most distal right-arm accessory region (4 CDSs), was rightwards transcribed.

The LSR integrase gene, named *int*/*rec*C2865-pp2, was detected at the left extremity of the phage (Fig 7). Two *int/rec*C2865-pp1-unrelated transcriptional regulators of the XRE superfamily (PF01381) were detected in front of *int/rec*C2865-pp2. Two 15-bp imperfect DRs (*attL* and *attR*) with consensus sequence 5’-[A/C]GG[A/T]GGAACGTTTGG-3’ were identified at the extremities of the phage integration core site, with 5’-aGGaGGAACGTTTGG-3’ (*attL*; which remained in the restored chromosomal bacterial gene upon prophage excision: *attB*) and 5’-cGGtGGAACGTTTGG-3’ (*attR*, carried in the excised circularized element: *attP*) allocated at each phage boundary (Fig 7). Comparative analysis of Int/RecC2865-pp2 revealed that eight *S. sciuri* genomes shared related integrases (Fig 7B). In all but one case we could determine that the integrated elements were incorporated truncating the same chromosomal gene as in MRSS C2865; however, no conservation of the DRs was observed in MRSS C2865 (Fig 7).

Unusually, an abortive infection bacteriophage resistance gene, designated *abiF* (Abi_2 family, PF07751), was detected at its accessory right-arm region. Proteins of this family are found in various bacterial species and contain sequences similar to the Abi group of proteins, which are involved in bacteriophage resistance mediated by abortive infection in *Lactococcus* species. AbiF shared highest identity (97% coverage, 58.1% ID) to the AbiF of food-associated biofilm-former *Staphylococcus cohnii* MF1844 (RefSeq assembly. No. GCF_001876725), among others.

#### C2865-pp3

C2865-pp3 was 126,192 bp and consisted of 166 CDSs of which only 34 (20.5%) had predicted functions. C2865-pp3 had a GC content of 30.8% and was integrated truncating a *yeeE/yedE* gene, coding for an inner membrane protein, positioned 819,965 bp downstream of *dnaA*. C2865-pp3 genome was organized into three proposed modules delimited by three different integrases (Fig 8). The leftmost 60.5-kb leftwards-transcribed arm carries genes whose predicted proteins are involved in lysogeny (integrase, DNA-binding protein), DNA metabolism (ribonucleoside reductase, DNA polymerase, deoxyribosyltransferase, methyltransferases, single strand DNA binding, DNA ligase, helicase DnaB) and adaption (Type II toxin-antitoxin system HicA and HicB). A centrally located 34-kb region, half leftwards and half rightwards transcribed, carried genes involved in lysogeny (integrase, represors), DNA replication (helicase), virion packaging (only TerL identified), and adaption (antirestriction ardA, Pro TGG-tRNA). The 31.5-kb rightmost rightward-transcribed phage genome region encompassed genes involved in phage morphogenesis (tape measure) and cell lysis (holin, endolysin) (Fig 8).

**Figure 8.**
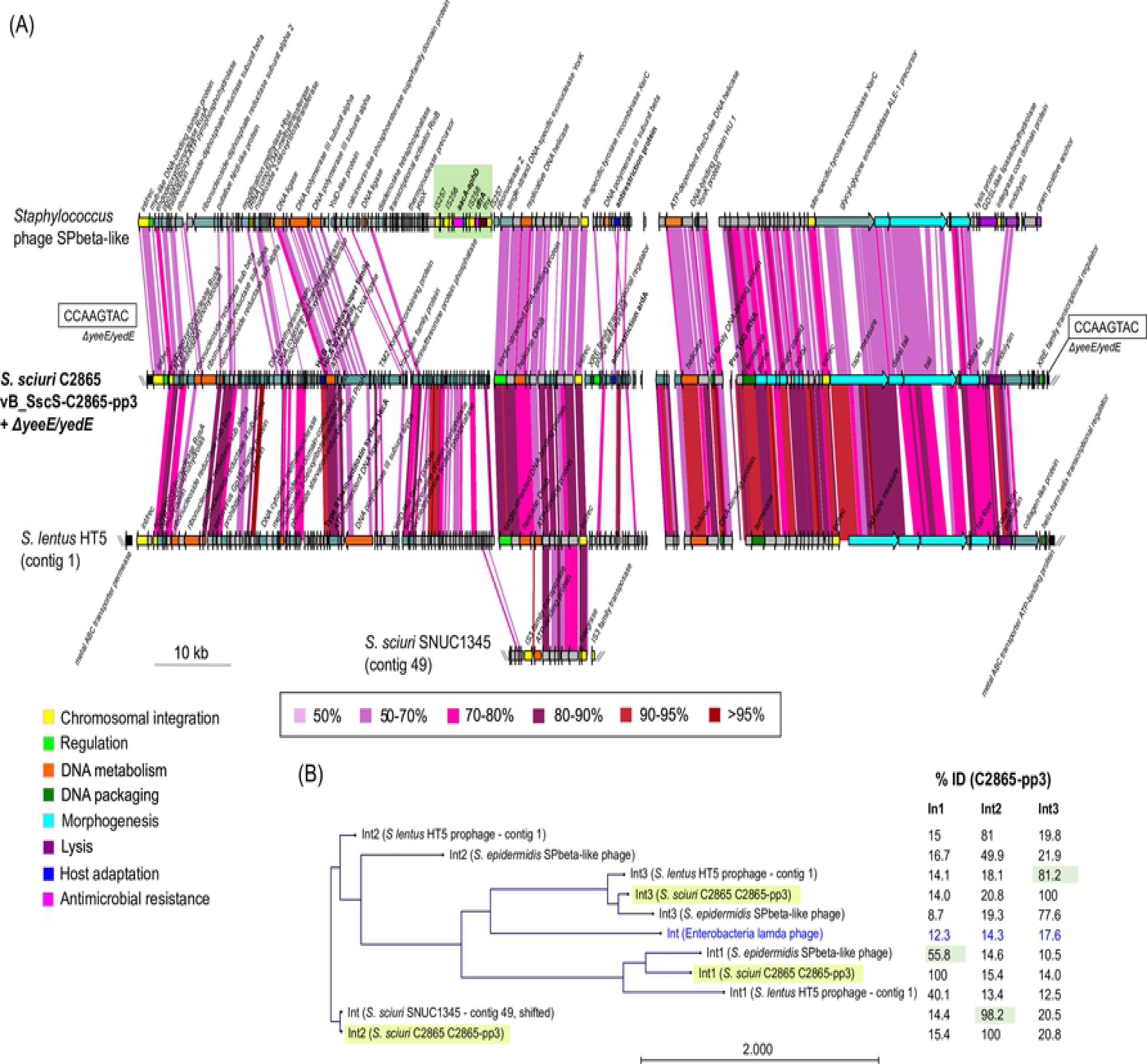
Graphical comparative analysis of prophage C2865-pp3 and closest relatives. (A) Graphical comparative analysis of the novel staphylococcal prophage vB_SscS-C2865-pp3 (C2865-pp3) integrated in the chromosome of *S. sciuri* strain C2865 and its closest relatives based on (i) all staphylococcal phages deposited in the Viral RefSeq database (accessed until November 2018) as well as on (ii) genetic elements enclosing the closest integrases in the NCBI pBLAST database (*Staphylococcus* phage SPbeta-like [Viral RefSeq acc. No. NC_029119]; *Staphylococcus lentus* HT5 – contig 1 – [RefSeq acc. No. NZ_SPOY01000001] and *S. sciuri* SNUC1345 – contig 49 – [NZ_QYJC01000049], respectively). Arrows denote the genes, length and orientation. Gene colors other than grey (hypothetical proteins) and grey-turquoise (others) represent genes involved in the following processes (color): chromosomal integration (yellow), regulation (green), DNA metabolism (orange), DNA packaging (dark green), phage morphogenesis (bright blue), cell lysis (purple), host adaptation (navy blue), antimicrobial resistance (pink). Areas of identity (tblastx, >50 aa match, >50% ID) between (pro)phages are indicated in colour scale. *Staphylococcus* phage SPbeta-like was linearized at position of interest. Area with green background denotes the region enclosing an independent mobile genetic region (flanked by two IS*257* copies in the same orientation), which encompasses the transposon Tn*4001* (IS*256*-*aacA/aphD*-IS*256*) as well as the trimethoprim resistance gene *dfrA*. (B) Maximum likelihood phylogenetic tree of the three integrases detected in C2865-pp3, as well as those present in its two closest (pro)phages (pBLAST search) (*Staphylococcus* phage SPbeta-like and *Staphylococcus lentus* HT5 – contig 1), and closest integrase (>80% amino acid identity, *S. sciuri* SNUC1345 – contig 49) and integrase of Enterobacteria lambda phage (NC_001416), used as outgroup (colored in blue). Integrases from C2865-pp3 are highlighted in faint yellow. Percentage of identity of all integrases analysed is indicated, with that sharing highest identity to respective integrase highlighted in faint green.

The leftmost located integrase gene, named *int1/rec*C2865-pp3, belonged to the LSR subfamily and was responsible for C2865-pp3 integration within *yeeE/yedE*. This integrase was preceded by a CDSs enclosing a DNA-binding domain of the of the transposase family ISL3 (PF13542), in the same orientation, which could behave as a transcriptional regulator. The centrally located integrase, named *int2/rec*C2865-pp3, belonged to the Y-rec family and was preceded by a XRE superfamily transcriptional regulator (PF01381) and by a phage antirepresor with two distinct antirepresor domains (PF02498, PF03374). The third integrase copy, located at the right arm and designated *int3/rec*C2865-pp3, also belonged to the Y-rec family; however, it lacked an discernible transcriptional regulator at its immediate surroundings. Instead, a CDS coding for a XRE superfamily transcriptional regulator was detected by CD-search (HTH_XRE, cd00093) 19 CDSs after *int3/rec*C2865-pp3, in the same orientation. The integration of C2865-pp3 generated two 8-bp perfect DRs (5’-GTACTTGG-3’) at its boundaries (Fig 8). Int1/RecC2865-pp3 and Int3/RecC2865-pp3 were unique in the entire BLASTp NCBI database. Instead, two hits were retrieved for Int2/RecC2865-pp3, one of them in the genome of *S. lentus* strain HT5 in a similar prophage-like region.

Two elements involved in phage adaption to the host were detected at its left and central regions, respectively: *hicA-hicB* (starting at position 23,791 of phage genome) and *ardA* (64,793). The *hicA* gene coded for an addition module toxin of the Type II toxin-antitoxin system (HicA_toxin family, PF07927). HicA_toxin is a bacterial family of toxins that act as mRNA interferases, while the antitoxin that neutralizes this family is HicB (Pfam domain PF15919). Immediately downstream of *hicA*, a hypothetical protein encompassing a HicB_lk_antitox superfamily domain (PF15919) was observed. Interestingly, an antitoxin *hicB* gene copy of the same superfamily was also identified at a different region of the *S. sciuri* C2865 chromosome (602,665 bp downstream of *dnaA*). The *ardA* gene coded for a putative antirestriction protein ArdA with an N-terminal domain (PF07275) involved in evasion of the bacterial Type I restriction modification system.

Moreover, C2865-pp3 harbored a transcriptional RNA with a Proline anticodon (Pro TGG tRNA) within the central region (phage position 81,705), which shared 86.7% nucleotide identity with corresponding tRNA of a number of Listeria phages (phage LP-124, LP-083-2, LP-064 and LP-125). The three prophages and the entire MRSS C2865 bacterial genome were then evaluated for codon usage and amino acid content to appraise whether acquisition of this tRNA could be an adaptive feature to favor phage replication, consequence of an increase TGG or proline within phage genome. However, proline was less abundant in C2865-pp3 (2.1%) than in MRSS C2865 genome (3.1%) and the other two phages (2.8-3%) (S7 Fig). TGG codon usage was alike in both C2865-pp3 and C2865 genome (0.85% each), while revealed increased in C2865-pp1 and C2865-pp2 (1.18, 1.17%, respectively) (S7 Fig).

C2865-pp3 shared closest identity to prophage-like element of mouse *S. lentus* HT5 and to *Staphylococcus* phage SPbeta-like, obtained from a clinical *S. epidermidis* 36-1. Remarkably, both phages also carried three different integrases (Fig 8). As expected by the low amino acid identity of the recombinase leading integration (40.1%), HT5-prophage-like element was incorporated at a different location, i.e. at the intergenic region of an ABC transporter ATP-binding protein and a ABC transporter permease genes (Fig 8), both present in the chromosomal DNA of *S. sciuri* C2865. Importantly, the left-arm region of SPbeta-like phage enclosed a composite transposon-like element (IS*257*-flanked) carrying a Dfr gene *dfrA* and transposon Tn*4001* (IS*256*-*aacA/aphD*-IS*256*), which harbors the aminoglycoside resistance gene *aacA-aphD*.

### ANI, phage classification and presumed packaging strategy

C2865-pp1 and C2865-pp2 shared highest identity each other (82% ANI; NCBI blastn 33% coverage, 88.3% ID). (S8 Fig). Alternatively, *Staphylococcus* siphophage SPbeta-like shared closest identity with C2865-pp3 (66% ANI; NCBI blastn 46% coverage, 72.91% ID) (S8 Fig) [58]. According to a recent reclassification scheme by Oliveira *et al.,* [58], based on phage genome homology, genome size and gene synteny, modular organization, percentage of functionally annotated predicted genes, temperate lifestyle and lack of or simple baseplate region, the three prophages were classified as S*iphoviridae*: C2865-pp1 and C2865-pp2 belonged to the staphylococcal phage cluster B, while C2865-pp3 clustered with SPbeta-like Shipophage, constituting the unique singleton so far among staphylococcal phages. Regardless of its large genome (126 kb), phage C2865-pp3 seems to belong to the *Siphoviridae* family as its putative simple base plate region (structural region highly conserved in *Siphoviridae*) shares high structural homology to that of lactococcal temperate *Siphoviridae* phage TP901-1, a model for *Siphoviridae* virion assembly [59].

According to TerL phylogenetic tree, C2865-pp1 and C2865-pp2 TerL clustered with typical *cos*-packaging phages, such as *S. aureus* 2638A phage. Hence, the predicted packaging mechanism used by these two novel phages seems to be mediated by cohesive ends [60] (S9 Fig). This was correlated in both phages with the presence of an HNH endonuclease gene in front of *terS*, specifically involved in DNA packaging of *cos* phages [61]. No packaging mechanism could be proposed for C2865-pp3, as its TerL was allocated distant from *cos/pac* characteristic clusters.

### The novel *S. sciuri* pathogenicity island (SscPIC2865) belongs to the SaPI4 family and related elements revealed common among *S. sciuri* genomes

A novel phage-inducible chromosomal island (PICI) of 13,101 bp, named SscPIC2865, was detected 20,705 bp upstream of *dnaA* gene. SscPIC2865 was classified as a *S. sciuri* pathogenicity island (SscPI) as it contains homologues of all core set of genes characteristic of *S. aureus* pathogenicity islands (SaPIs) and displays synteny with previously characterized SaPIs. SscPIC2865 consisted of 21 CDSs, of which nine (42.9%) lacked any predicted function (Fig 9A). SscPIC2865 contained a site-specific integrase gene (*int/rec*) of the Y-rec family at its left arm end, required for the integration of the element. SscPIC2865 Int/Rec directed chromosomal integration at the end of the 3’ end of the 30S ribosomal protein S18 gene (*rpsR*). Two SscPIC2865-flanking 15-bp DRs (5’-AAAGAAGAACAATAA-3’), constituting the attachment core sites of SscPIC2865 (*attL*, *attR*) recognised by Int/Rec, were detected (Fig 9A).

**Figure 9.**
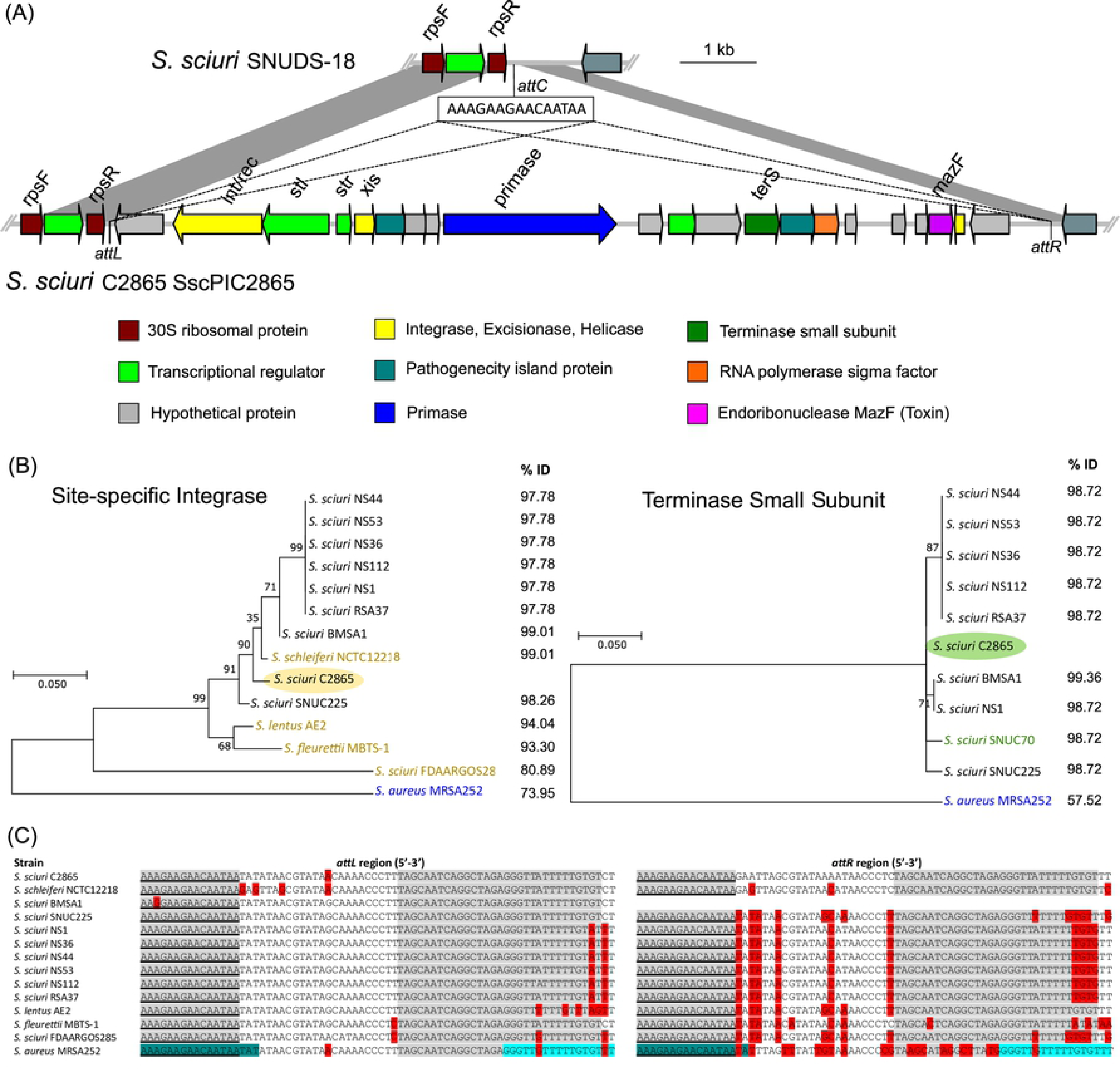
Novel staphylococcal pathogenicity island SscPIC2865 and analysis of integrases, terminases and integration sites of related elements. (A) Schematic representation of novel *S. sciuri* C2865 staphylococcal pathogenicity island, named SscPIC2865, plus the integration immediate flanking region in the chromosome, downstream the 30S ribosomal S18 protein (*rpsR* gene), and comparison of the integration region with that of *S. sciuri* SNUDS-18. Arrows denote the genes, length and orientation. The core integration site sequences detected (5’-AAAGAAGAACAATAA-3’) as well as position for integration (*attC* for putative integration site in *S. sciuri* SNUDS-18 chromosome; *attL* and *attR* at both extremities of SsPIC2865) are indicated. Selected genes and resultant protein functions are as follows (5’ to 3’ read out) (color): *int*, integrase (yellow); *stl*, repressor of leftward SsPIC2865 transcription (green); *str*, repressor of rightward SsPIC2865 transcription (green)*; xis*, excisionase (yellow); *pri*, primase (navy blue); putative transcriptional regulator (green); *terS*, terminase small subunit (dark green); RNA polymerase sigma factor (orange); *mazF*, endoribonuclease MazF – toxin of toxin-antitoxin system) (purple); and helicase (yellow). Areas of nucleotide similarity (nblastn, >100 bp match, >99% ID) between both structures are indicated in grey. (B) Molecular Phylogenetic analysis by Maximum Likelihood method of amino-acid sequences of (left) all site-specific integrases (Int) deposited in the protein NCBI database and (right) all terminases small subunit (TerS) sharing >80% amino acid identity with respect to those present in SscPIC2865 (*S. sciuri* NS44 [NCBI RefSeq. No. NZ_LDTO01000002]; *S. sciuri* NS53 [NZ_LDTP01000017]; *S. sciuri* NS36 [NZ_LDTN01000005]; *S. sciuri* NS112 [NZ_LDTL01000018]; *S. sciuri* NS1 [NZ_LDTK01000052]; *S. sciuri* RSA37 [NZ_LDTQ01000004]; *S. aureus* BMSA1 [NZ_LFXB01000008]; *S. sciuri* SNUC225 [NZ_PZHF01000049] for both Int and TerS; *S. schleiferi* NCTC12218 [NZ_RXWV01000003]; *S. lentus* AE2 [NZ_SPPP01000007]; *S. fleurettii* MBTS-1 [NZ_MWJM01000005] and *S. sciuri* FDAARGOS285 [NZ_CP022046] for Int; and *S. sciuri* SNUC70 [NZ_PZHH01000006) for TerS), in addition to those of prototype SaPI4 from *S. aureus* MRSA252 (GenBank acc. No. BX571856), which integrase belongs to *att*/integrase family I [62] (displayed in blue). Strains with Int but either lack of or divergent (<50% ID) TerS, and vice versa, are indicated in yellow or green color, respectively. Percentage of amino acid identity with respect to Int or TerS of SscPIC2865 are displayed on the right of the respective tree. TerS-carrying strains corresponded to those harboring the SscPIC2865-conserved integrase in all cases but one (*S. sciuri* SNUC70), which was integrated at a different position of the bacterial chromosome (at tRNA, *ssrA* gene). (C) Sequence comparison analysis of the phage-related chromosomal island *att* sequence core site region (*attL* and *attR*) of SscPIC2865 and corresponding region of all strains carrying a similar integrase (>80% ID), in addition to that of *S. aureus* MRSA252 (harboring prototype SaPI4), which also integrates at the 3’-end of the S18 ribosomal gene *rpsR*. Bases in grey background indicate perfect matches between upstream and downstream integration region among all included strains. Underlined area represents the proposed integration core sequence of SscPIC2865 and related elements. Bases differing from the *att* region of *S. sciuri* C2865 SscPIC2865 are indicated in red. Original *att* core site length of SaPI4 is indicated in dark blue [62], and the additional direct repeat found is shown in blue.

Immediately upstream of this *int/rec* gene, two divergently oriented CDSs encoding DNA-binding proteins of the HTH_XRE superfamily (CD-search cl22854), termed Stl and Str in elements of the SaPI family, were detected [62]. Stl is essential for maintenance of lysogeny while Str is responsible for activation of the lytic cycle prior mobilization by a helper phage. An excisionase gene (*xis*), involved in excision of SaPI-like elements upon regulation of these repressors, was located downstream of *str* gene. In the middle of the island, a primase gene [D5 N-terminal like domain (PF08706), characteristic of DNA viruses and P4 DNA primase phages] fusioned with a replication initiation protein gene [C-terminal domain (PF032288)] was detected. This was coupled with an AT-rich iteron region at its 3’-end, comprising the putative replication origin. Only two hits >80% identity to SscPIC2865 primase were detected : primase of *S. fleurettii* MBTS-1 (96.5% ID, NCBI RefSeq. No. NZ_MWJM01000005) and that of *S. sciuri* SNUC740 (80.6% ID; NZ_PZHE01000007). Downstream of this area, a copy of a terminase small subunit (*terS*) of the family Terminase 2 was detected (Fig 9A). None of the genes involved in interference of the helper phage reproduction in favor of SscPIC2865 packaging were identified. This may be due to lack of sequence conservation, typical of SaPI-containing genes. However, this 3’ region carried a putative transcriptional regulator, a RNA polymerase sigma factor and a putative helicase, which might be involved in the replication of the element (Fig 9A). Interestingly, a putative *mazF* toxin gene, coding for an endoribonuclease (mRNA interferase) of a Type II t oxin-antitoxin system (MazF/MazE) was identified in the accessory region of SscPIc2865 by HHpred (9.3e-20 E-value).

Twelve integrases sharing >80% amino acid identity to SscPIc2865 integrase were identified by sequence comparison, all truncating the 3’-end of the ribosomal gene *rpsR*, (Fig 9B). Of these, nine belong to *S. sciuri* and the rest to *S. schleiferi*, *S. lentus* and *S. fleurettii*. Remarkably, their integrase-surrounding regions share the characteristic structure of SaPIs (data not shown).

Among the six groups of site-specific integrases detected so far among elements of the SaPI family, SaPIs of integrase group I, with SaPI4 as prototype, also integrate at the 3’ region of the ribosomal protein S18 (*rpsR* gene) [62]. However, the amino acid identity of SaPI4 (*S. aureus* MRSA252) and SscPIC2865 integrases was considerably low (74%) (Fig 9B). To address whether SscPIC2865 integrase and relatives belong the SaPI4 family, the integration region (*attL*, *attR*) of these elements was analyzed in further detail (Fig 9C). The SscPIC2865 perfect DR 5’-AAAGAAGAACAATAA-3’ (15 bp) was detected at both extremities of the islands in all cases, including in *S. aureus* MRSA252, except for *S. sciuri* strain BMSA1 (Fig 9C). Hence, we propose SscPIC2865-related elements belong to the same Int group I, with conserved core site 5’-AAAGAAGAACAATAA-3’, instead of the slightly longer formerly deduced attachment site of Int group I 5’-AAAGAAGAACAATAATAA-3’.

Comparative analysis of SscPIC2865 TerS, mostly conserved in SaPI-like elements, identified nine terminases sharing >80% identity to TerS of SscPIC2865 (Fig 9B). These corresponded to strains harboring the SscPIC2865-conserved integrase in all cases but one (*S. sciuri* SNUC70).

Sequencing analyses of potential CIs and re-storage of SscPIC2865 chromosomal integration site revealed that SscPIC2865 had the ability to excise the bacterial host genome reestablishing the expected *attP* and *attB*, in the genomic island and bacterial chromosome, respectively.

## Discussion

We determined the complete genome of a canine MDR MRSS strain from Nigeria and identified a novel staphylococcal TMP resistance gene (*dfrE*) located within a novel MDR mobilizable plasmid, which harbored additional adaptive traits. Our WGS approach allowed us to resolve its additional mobilome, unveiling high genome plasticity enriched in chromosomal and extrachromosomal novel mobile elements. Regardless the ubiquity of *S. sciuri* in different ecological niches and its proven role as reservoir for clinically relevant AMR genes, very limited WGS data are available in comparison with *S. aureus* or the CoNS *S. epidermidis*. In addition, most genome sequences are draft genomes generated by de novo assembly of short reads, which do not enable the complete determination of elements containing repeat regions, such as ISs. As observed here, ISs are highly abundant not only among AMR-carrying mobile elements, but also across the staphylococcal chromosome. Here, the combination of short-read Illumina sequencing followed by deep long-read PacBio sequencing using large DNA fragmentation allowed the determination of the complete genome and mobilome at the minimum mutation rate. Therefore, our sequencing approach evidenced the necessity for long-read sequencing to establish and complete the landscape of larger mobile elements surrounded by repeat regions. A total of 11.2% of *S. sciuri* C2865 genome consisted of the MGEs here described, which increased to over 13% when considering the length of the 78 predicted ISs (estimated IS length: 700 bp). Of note, all functionally proved AMR genes were carried by MGEs, evidencing HGT acquisition.

Phylogenomic analyses of the 30 *S. sciuri* group genomes analyzed revealed that MRSS C2865 clustered within a *S. sciuri* species sub-branch, which included three additional strains: two from human and animal infections (*S. sciuri* Z8, SNUD-18) - indicating pathogenicity potential at the genomic level - and one from food, *S. sciuri* CCUG39509. The latter was originally considered the type strain of *S. sciuri* subsp. *carnaticus* (ATCC 700058). Nevertheless, the subspecies division into *S. sciuri* subsp. *sciuri*, *S. sciuri* subsp. *carnaticus* and *S. sciuri* subsp. *rodentium* is currently rejected [4, 90]. Based on a range of phenotypic, biochemical, physiological and genetic analyses, Svec *et al.,* [4] recently showed high interspecies heterogeneity with no clear differentiation into the different subspecies. While agreeing with those statements, our WG-based phylogenomic analysis revealed a clear distinction of an intraspecies sub-cluster, sharing 96% ANI with the rest of the *S. sciuri* genomes. This ANI value falls within the threshold to be considered a different species (≈95–96%) [73]. Hence, we suggest that there indeed may be a subspecies discrimination among the *S. sciuri* species, but such distinction needs to be addressed by WGS comparisons and might not correspond to the outcomes retrieved by the abovementioned traditional methods. Alternatively, the available metadata of compared genomes evidence lack of clear phylogeny demarcation depending on the origin, source or host. This corroborates the low host tropism already suggested for this species [3]. The pan genome analysis of all 21 *S. sciuri* genomes included revealed that MRSS C2865 significantly exhibited the highest number of both total and unique genes, and that most of them (75%) corresponded to the MGEs identified. These values highlight the particularity of MRSS C2865 as acceptor of adaptive mobile traits and the importance to search for environmental niches to understand the evolution of this exceptionally versatile bacterial species. At the species level, the open pan genome identified, with an accessory gene content of over 73%, highlights an important source of evolutionary novelty that facilitates rapid adaptation via HGT. Hence, genome-wide association studies among this species and additional commensal staphylococci are needed to identify and understand their exchange platforms and mechanisms, important to track their genome evolution.

Importantly, the novel *dfrE* alone conferred high level TMP resistance in both staphylococci and *E. coli*. A potential environmental origin is predicted, as Dhfr from soil-associated *P. anaericanus* evidenced the closest relative and DfrE was remarkably distant from the typical staphylococcal TMP resistant Dfrs. Scarce data is available on *P. anaericanus*, but bacteria belonging to this genus are ubiquitous in nature, and closest species have been reported in different environmental sources, such a soils and rhizosphere from different crops [91–93]. We identified the *dfrE* gene in a limited number of staphylococcal genomes from *S. sciuri*, *S. aureus* and *S. arlettae* of human and animal origin (including clinical samples) as well as in animal-associated *M. caseolyticus* strain JCSC5402 [47] and *Exiguobacterium* sp. strain S3-2 [46]. However, only in the later strain the *dfrE* gene, which authors denominated *dfr_like*, was denoted as AMR gene and proved to confer TMP resistance in *E. coli*. *Exiguobacterium* spp. are extremophiles adapted to a wide range of habitats, including cold environments. They are also recovered from closer human-associated niches, such as aquaculture, landfills, plant-derived foods, food-processing plants and pharmaceutical wastewaters [94, 95], including AMR strains [46, 96, 97]. These data evidence the transferability of clinically relevant AMR genes across diverse bacteria from different taxonomic families (order Bacillales) from diverse environments. All *dfrE*-carrying genomes harbored it in MDR plasmids or plasmid-associated elements, hosting additional antibiotic and /or metal resistance genes. This co-localization is of concern, because it enables the transfer of diverse adaptive traits via a single HGT event. Importantly, in *Exiguobacterium* sp. strain S3-2, the *dfrE* gene, enclosed within plasmid pMC2, was enclosed within an integral Tn*3*-like transposon [46]. This element was flanked by two typical 38-bp IRs involved in excision and integration, common in this transposon family members. pUR2865-34 harbored a truncated version of this element, as only the IS*1216E-res/rec2*-*res/rec3-dfrE-thy* region (4,345 bp), including the 3’-end 38-bp IR, was conserved. Indeed, this region was highly preserved in all additional *dfrE*-carrying strains but *S. sciuri* GN5-1 pSS-04. Here, the *dfrE*-carrying region seems to have evolved afterwards by losing the *res/rec2* plus two immediate downstream genes from the Tn*3*-like remnant. This indicates that *Exiguobacterium* sp. harboured an ancestral *dfrE* (*dfr_like*) transposable element that has recombined and jumped to different staphylococcal species and to macrococci from a single ancestor. The *ica*-locus variant detected within pUR2865-34 did not prompt remarkable biofilm formation when tested in the CV assay, as formerly observed for its closest relative in pAFS11 in MRSA Rd11 [44]. However, *ica*-locus variant carrying strains tested positive to the colorimetric test. Its evolutionary history and *in-vivo* activity will be the object of further characterization.

The backbone of mosaic pUR2865-34 denoted a theta replication mechanism, characteristic of larger staphylococcal plasmids. The N- and C-terminal domains of the encoded RepA were homologous to different RepA_N proteins, reflecting the ability of these proteins to undergo domain replacements, which are presumably driven by selection pressures and host/plasmid incompatibility [98]. The putative Type Ib partitioning system detected in pUR2865-34 is well studied in Gram-negative bacteria, but little to nothing in Gram-positive cocci. Nevertheless, these systems contribute to the prevalence and spread of these plasmids, ensuring stable inheritance and effectively maintain resistance in the absence of selection. Although this plasmid was not conjugative, it contained a mimic of an origin of conjugative transfer, very similar to those recently identified by Bukowski *et al.,* [45]. Hence, pUR2865-34 could only move by conjugation via this *oriT* mimic sequence in the presence of a pWBG749-like plasmid, or via the integrated *mob* gene and associated *oriT* of pUR2865-int in the presence of a conjugative plasmid [43]. However, MRSS C2865 did not harbor any member of the three known staphylococcal conjugative plasmid families (pSK41, pWBG4 and pWBG749) nor any recognizable transfer gene cluster that could act *in-trans.* Indeed, conjugative plasmids are rare in staphylococci, representing only ca. 5% of plasmids in *S. aureus* [99].

The novel MDR SCC*mec*C2865 enclosed additional AMR genes, remarking an IS*1216E*-flanked region carrying the tetracycline resistance gene *tet*(S), which IS copies may have facilitated its capture. This gene has been found among *Firmicutes* and Gammaproteobacteria from diverse ecological sources since 1950s [100]. However, the *tet*(S) has only been detected twice before in staphylococci: (i) among MRSA isolates from animal carcasses [101] and (ii) within the *S. sciuri* SCC*mec*C2865-related SCC*mec*TG24 cassette, from ready-to-eat meat [15]. The additional closest cassette corresponded to *mecA-mecC* hybrid SCC*mec*-*mecC* in *S. sciuri* GVGS2 from a bovine infection [23]. These strain sources highlight that *S. sciuri* from animals behave as reservoirs for mosaic MGEs carrying AMR genes from different species. CcrA/B recombinases mediate SCC*mec* excision, circularization, site-specific chromosomal integration and recombination reactions between SCC*mec att*-flanked regions. Unusually, SCC*mec*C2865 lacked any recognizable *ccr*-encoded recombinase. The ancestral Ccrs might have been removed by IS flanking deletions rendering this cassette not mobile; however, no deleterious regions flanking the cassette were observed. Alternatively, all SCC*mec* Ccr enclose a conserved motif that was only detected in the LSR of two novel prophages C2865-pp1 and C2865-pp2, and shared closest phylogenetic distance to CcrC allotypes. Hence, even though neither circular intermediates nor an intact *rlmH* gene could be detected using the primer sets designed, the possibility that these prophage-located LSR participate in any of the abovementioned functions is tempting. Of note, virtually all staphylococci contain prophages and the transduction of small SCC*mec* types and SCC*mec* internal modules between compatible *S. aureus* strains has been demonstrated [30, 33, 34].

We detected for the first time in *S. sciuri* the integration of transposon ψTn*554* at the chromosomal *radC* gene. Further insights revealed that several *S. sciuri* genomes exhibited this *radC* gene disrupted, always by the presence of a transposon-like element - initially detected among enterococci, which harbors an AMR gene cluster carrying the emerging oxazolidinone and phenicol resistance gene *optrA* and the phenicol resistance gene *fexA* [102]. This observation signals the *radC* gene of *S. sciuri* as hotspot for integration and recombination of mobile adaptive elements coming from different bacterial backgrounds.

A novel chromosome excisable PICI, named SscPIC2865, was identified. PICIs are characterized by a specific set of phage-related functions that enable them to hijack the phage lytic reproduction cycle of helper phages for their own high-efficient transduction. These elements normally carry critical staphylococcal virulence genes, such as Panton-Valentine Leukocidin, toxic-shock syndrome, enterotoxins and other superantigens [62]. However, some PICIs do not carry any noticeable pathogenicity-related accessory genes. Unlike other members of the SaPI4 family, SscPIC2865 harbored a putative *mazF* toxin gene of a toxin-antitoxin system type II (*mazEF* locus) in its accessory region. MazF is a sequence-specific RNase that cleaves several transcripts, including those encoding pathogenicity factors. In addition, several staphylococcal strains from WGS projects harbored related elements at the same integration site. Particularly, *S. lentus* AE2, *S. fleurettii* MBTS-1 and *S. schleiferi* NCTC12218, either lacked an identifiable TerS or harbored a highly diverse TerS. In addition, *S. sciuri* strain BMSA-1 harboured a 3-‘end mosaic structure carrying a 23S rRNA methyltransferase Erm gene sharing 78.5% nucleotide identity with the recently described MLSB resistance gene *erm*(44)v [103]. These observations evidence that SscPIC2865-related elements are common among *S. sciuri* and have counterparts or remnants in different staphylococcal species, contributing to genome plasticity and potential toxigenicity. Hence, the ribosomal protein S18 (*rpsR* gene) appears as a hub for integration of mobile islands and should be consider when addressing staphylococcal chromosomal-integrated elements.

MRSS C2865 resulted to be a polylysogen, with three novel unrelated prophages, which accounted for 7.2% of the bacterial genome length. Despite the ecological importance of *S. sciuri* within the staphylococcal genus, only two former reports have identified *S. sciuri* phages: (i) three myoviruses from urban sewage in Poland [26], and (ii) two temperate siphophages from UV-induced *S. sciuri* strains [27], which were highly different from our novel phages. C2865-pp2 shared homology with two putative *S. sciuri* prophages; however, only C2865-pp2 enclosed a predicted adaptive gene at its accessory region: a bacterial abortive infection protein (AbiF). These systems are considered as “altruistic” cell death mechanisms that are activated by phage infection and limit viral replication, thereby providing protection to the bacterial population [104]. Hence, resultant AbiF might be involved in superinfection immunity, to prevent the infection of similar phages at the bacterial population level. As recently observed by Oliveira *et al.,* [58], a low rate of staphylococcal phages may exhibit a variety and uncommon number of site-specific recombinases, as it was the case for phage C2865-pp3, which enclosed 3 different integrases. Phage integrases are required for the establishment of the lysogeny, but its maintenance is also dependent on the regulatory proteins next to it [58]. At least two of these three integrases were surrounded by transcriptional regulators and/or antirepresor proteins. This may enable the integration at additional chromosomal sites depending on the integrase used. In fact, the closest putative prophage enclosed within animal *S. lentus* HT5, also carrying three different integrases, was integrated at a different location. According to a recent staphylococcal phage re-classification scheme [58], C2865-pp3 falls within the *S. epidermidis* SPbeta-like singleton, which also encloses three integrases but was reported to lack genes associated with stable lysogeny maintenance. Here, we show that a highly related phage can indeed stably reside as a prophage. Importantly, C2865-pp3 enclosed several adaptive genes, highlighting the presence of a putative type II toxin-antitoxin system of the HicA family (HicA/HicB) and an antirestriction protein (ArdA family). These elements could promote maintenance of the phage and/or avoid cell damage as well as to evade restriction in the recipient bacterium by the host restriction enzyme systems, respectively. Finally, the role of the tRNA unique to C2865-pp3, which shared homology to that present in *Listeria* spp., remains elusive, as its presence was not justified by an increased in the specific codon or amino acid usage [105]. Importantly, SPbeta-like phage harbored an AMR gene cluster enclosed within a composite transposon-like element, harboring resistance determinants for aminoglycosides and TMP. This is a central feature demonstrating that C2865-pp3-related functional phages can indeed behave as reservoirs for AMR genes, which can be further spread into different bacterial hosts. Whether the different lysogenic modules of these phages are functional and widen their infectivity range and site-integration versatility still needs to be clarified, but the fact that closest relatives were retrieved from different bacterial species (*S. epidermidis*, *S. lentus*) strongly points towards this possibility. In addition, the enclosed adaptive features detected reflect the impact of staphylococcal prophages in genome evolvability.

The conservation of the integrase-generated DRs flanking the three prophage genomes, together with their ability to excise the bacterial chromosomal DNA and circularize, as well as the high genome synteny observed among unrelated (pro)phages strongly suggest that these prophages are functional. If this is the case, along with progeny generation, also plasmid s or chromosomal DNA of the bacterial host may be mistakenly encapsulated [29]. Although it is generally assumed that this process is mainly performed by *pac* phages (headful packaging mechanism), the ability of some *cos* phages – which are more sequence-specific - to pack determined PICIs has been also proved [61]. Indeed, phage-mediated HGT is one of the primary driving forces of bacterial evolution. However, transduction of AMR genes is poorly understood. Hence, it is conceivable that any of these prophages may package and mobilize any of the MGEs discovered here. This hypothesis is supported by the fact that no conjugative element that could mediate mobilization of any of the three plasmids was present. These prophages could also contribute to further genome evolution by mediating the mobilization of the novel *ccr*-lacking SCC*mec* element and SscPIC2865 islands. In fact, PICIs of the SaPI4 family are only induced by endogenous prophages [62]. Further studies are warranted to explore the transduction ability of these prophages to further spread the novel MGEs identified.

## Conclusions

Our work unveils a number of novel mobile elements, including a novel environment-associated staphylococcal TMP resistance gene already spread in different bacterial species with different epidemiological backgrounds. A novel MDR mobilizable plasmid, SCC*mec*, PICI and three unrelated prophages are also uncovered here. We confirm the ubiquity, high genome plasticity and low host tropism of *S. sciuri* highlighting its role as a resourceful reservoir for evolutionary novel features contributing to its extraordinary versatility and adaptability. Monitoring the mobilization platforms for spread of this novel MDR resistance plasmid and additional mobile elements is critical to understand and refrain further expansion of AMR. Hence, upcoming studies will determine the mobilization ability of these novel MGEs and the role of the enclosed novel prophages to mediate such events. We reassure and stress the importance of refined genome-wide ecological studies to understand and eventually refrain the recombination and mobilization landscape of this critical opportunistic pathogen. We also highlight the need to profit from the richness of genomic data so far available in combination with functional analyses to unveil adaptive traits that otherwise will pass unnoticed among the overwhelming sequencing scenery.

## Materials and methods

### Bacterial strain selection and characteristics

MRSS strains C2865, C2853, C2854 and C2855 were obtained in a former study on the occurrence of MRCoNS from canine samples, and were recovered from the groin area of unrelated stranded dogs in Nskuka, Nigeria [13]. These strains exhibited TMP resistance while none of the known staphylococcal TMP resistance genes (*dfrS1*, *dfrD*, *dfrG*, *dfrK*) [35, 36, 38, 39], nor the streptococci/enterococci *dfrF* gene [37, 40] were detected. All strains were MDR and exhibited related AMR patterns. Strain C2865 was selected for WGS in order to unveil the genetic basis for TMP resistance.

### DNA extraction, whole-genome sequencing, assembly and annotation

Detailed DNA extraction procedures are described in S1 Data. Briefly, genomic DNA was extracted by two different methods: for Illumina sequencing, the Wizard Genomic DNA Purification Kit was used including both lysozyme and lysostaphin (10 mg/ml each) (Ref. A1120, Promega Corporation, Spain); PacBio sequencing, DNA was isolated using a phenol-chloroform method with some modifications for improved cell lysis. Plasmid DNA was obtained using the GenElute Plasmid Miniprep Kit (Ref. PLN350, Sigma) also including a lysis step with lysozyme (2.5 mg/ml) and lysostaphin (0.25 mg/ml) for 20 min at 37°C after the resuspension solution step.

High-throughput WGS of *S. sciuri* C2865 DNA was performed with Illumina Miseq (2 × 300 bp), with NEBNext Ultra kit (http://www.genomicsbasel.ethz.ch). For long-read sequencing, PacBio RSII was used prior DNA fragmentation of 15 kb followed by a mild size selection. Illumina read quality was checked by Fastqc (https://www.bioinformatics.babraham.ac.uk/projects/fastqc/) and reads were trimmed using Trimmomatic v0.36 [63]. Good-quality Illumina reads were de-novo assembled using SPAdes [64]. PacBio RSII raw reads were assembled using Canu [65]. Protein-coding genes, tRNAs and rRNA operons were predicted using Prodigal [66], tRNAscan-SE and RNAmmer [67]. Predicted protein sequences were compared against the NCBI nr database using DIAMOND [68], and against COG [69] and TIGFRAM [70] using HMMscan [71] for taxonomic and functional annotation. Genomic alignment dot-plots between Illumina and PacBio resulting contigs were generated with D-GENIES software to evaluate consistency and reliability of both sequencing and assembly approaches [72]. Amino acid and codon usages were determined for C2865 chromosome and prophages using the compareM package (https://github.com/dparks1134/CompareM).

### *S. sciuri* group phylogenomic analysis, comparative genomics and pan genome examination

All genomes available belonging to the *S. sciuri* group species were downloaded from the NCBI database (accessed until April 2018). In total, thirty strains (29 from NCBI, MRSS C2865) were included, which belonged to the following species (no of strains): *S. sciuri* (21), *S. lentus* (5), *S. vitulinus* (2), *S. fleuretti* (1) and *S. stepanovicii* (1) (S1 Data). In order to measure the probability of two genomes belonging to the same species, an ANI of the 30 *S. sciuri* group genomes was calculated using JSpecies as indicated before [73]. A heat map was generated using the ANI matrix output table with R [74]. In parallel, a maximum likelihood tree for all the *S. sciuri* group genomes was generated using RAxML (version 7.2.6) [75] using core alignment obtain with Parsnp software within Harvest Suite package [76]. The results were visualized using CLC Genomics Workbench 12. BLAST Ring Image Generator (BRIG) was used to evaluate and visualize comparisons between MRSS strain C2865 and its closest genomes (>98% ANI), using MRSS C2865 genome as reference [77]. Pan genome analysis (core plus accessory genome) for the 21 *S. sciuri* genomes was carried out using Roary with a 95% identity cutoff value [78].

### Detection and analysis of resistome and mobilome from *S. sciuri* C2865

#### Antimicrobial resistance (AMR) genes

AMR genes formerly detected in MRSS C2865 (S1 Table) were blasted against the WGS of strain C2865. Contig/s were manually checked for redundant *dhfr* genes and for additional resistance genes of interest (S1 Data for Illumina data).

#### Plasmids

Plasmid contig identification and plasmid reconstruction was achieved by contig coverage, sequence similarity with plasmid backbone genes, gene composition and organization and contig boundaries redundancy (circularity) (S1 Data for Illumina data). Putative *oriT* and *oriT* mimics on the mobilizable elements were searched using the core sequence of those from conjugative plasmids [79]. For RCR plasmids, *dso* and *sso* regions, involved in the initiation of replication of the leading and lagging strand, respectively, were searched using core sequences of a RCR representative per plasmid family [80]. The secondary structures of *dso, sso, oriT* and *oriT* mimic were generated using Mfold web server for single-stranded linear DNA at default parameters [81].

#### Prophages

Detailed analyses are indicated in S1 Data. Manual inspection of phage-associated genes (morphology/structure, lysogeny, cell lysis, DNA metabolism) and characteristic functional modular organization was implemented for phage confirmation and integrity. Integrase motif and domain analysis of the translated candidate CDSs was performed against ScanProsite database [82], Pfam database [83], and NCBI conserved domain database (CDD) [84]. Integrase-directed generation of DRs as a result of genome integration was investigated manually by sequence comparison of bacterial chromosome-prophage boundaries with corresponding regions of MRSS C2865 and a *S. sciuri* strain prototype (SNUDS-18, Genbank ac. No. CP020377) lacking those prophages. ANI pairwise comparison between C2865-enclosed prophages and all staphylococcal phage genomes available in the Viral RefSeq database (accessed until November 2018, n=187) were calculated using the JSpecies with default parameters [73]. A heat map was generated using the ANI matrix output table with R [74].

#### Staphylococcal Chromosomal Cassette *mecA* (SCC*mec*)

ORFs found downstream of the integration gene 23S rRNA (pseudouridine(1915)-N(3))-methyltransferase RlmH, initially known as OrfX, as well as regions containing characteristic SCC*mec* genes (*mecA, mecR, mecI, ccr*) were analyzed. To detect *ccr* gene/s, which resultant proteins belong to the large serine recombinase (S-rec) (LSR) family, consensus motif Y-[LIVAC]-R-[VA]-S-[ST]-x (2)-Q derived from Prosite entry PS00397 (http://prosite.expasy.org) was used (S1 Data). Additional S-rec encompassing the Y-[LIVAC]-R-[VA]-S-[ST]-x (4)-Q motif were likewise screened along the entire MRSS C2865 genome. Putative ISS for SCC*mec* or *att* core sites recognized by the typical staphylococcal Ccr were manually identified by searching for the consensus sequence 5’-GAAGC[AG]TATCA[TC]AAAT[AG]A-3’ (S1 Data).

#### Others

Additional recombinases associated with MGEs were investigated as above for motif and domain analysis in translated gene candidates of MGEs (S1 Data). SscPI chromosomal integration site was compared with similar integrase-carrying genomic islands (S1 Data). Integrase-directed generation of DRs as a result of SscPI integration were investigated by sequence comparison of bacterial chromosome-SscPI boundaries with corresponding regions of *S. sciuri* strain SNUDS-18, which lacks any insertion in corresponding chromosomal region. In addition, ISsaga2 web tool was used for IS identification and quantification [85].

#### Excision ability of chromosomally located MGEs

Potential excision and circularization of selected chromosomally located mobile elements, in addition to detection of resultant chromosomal region after excision were tested by specific inverse and conventional PCR, respectively (S1 Data, S1 Table).

#### Phylogenetic analyses of proteins of interest

Phylogenetic analyses for Dhfr/Dfr and Ccr of interest were investigated by the construction of Neighbor-joining tree using the MEGA 7.0.21 program (S1 Data) [86]. All site-specific tyrosine integrases (Y-int) and terminases small subunit (TerS) in the NCBIp database sharing >80% amino acid identity to those of SscPI in C2865 were correspondingly analyzed.

A maximum likelihood phylogenetic tree for MRSS C2865 prophage integrases and all related integrases was built upon amino acid sequence alignment as above indicated (CLC Genomics Workbench 11) (S1 Data). To estimate the packaging mechanism of identified prophages, a circular phylogenetic tree of the terminase large subunit (TerL) from MRSS C2865 prophages and those identified in the 187 staphylococcal phages deposited in the Viral RefSeq database was likewise created upon amino acid sequence alignment using CLUSTALW (CLC Genomics Workbench 11). TerL of staphylococcal phages with known packaging mechanism was indicated [60].

#### Construction of recombinant plasmids for DfrE functionality, transfer assays and plasmid integrity

To address functionality of the candidate TMP resistance gene and for potential synergistic activity by its immediate 3’-end thymidylate synthase gene (*thy*), three different *dfrE* containing regions were amplified and cloned into pBUS-Pcap-HC (constitutive promoter) or pBUS-HC vectors using the Gibson assembly workflow: (i) *dfrE* gene alone, (ii) *dfrE* gene plus flanking regions, and (iii) *dfrE* gene, *thy* gene and flanking regions (Table S2; S1 Data) [50]. Constructs of interest were transformed into *Escherichia coli* DH5α and subsequently into *S. aureus* RN4220 (S1 Data).

Purified DNA of two independent transformants containing the original *dfrE* gene (RN4220/pUR2865-34) were sent for Illumina WGS (HiSeq, 10 Mill reads, 2 x 150 bp) for confirmation of integrity of transformed plasmid. In parallel, one microgram of *dfrE*-containing plasmid DNA from selected transformants (RN4220/pUR2865-34) and from original strains (C2865, C2853, C2854, C2855) were digested with 1 Unit of S1 enzyme (1 min at 37°C) (Thermo Fisher Scientific) for plasmid linearization, and plasmid length and integrity was assayed by gel electrophoresis.

#### Antimicrobial susceptibility testing

MIC of TMP was determined in strains of interest by the agar dilution method on Mueller-Hinton plates (MH, Becton Dickinson) with a concentration range of 0.5 to 4’096 µg/ml (S1 Data) [87]. The agar disk-diffusion method was also used to test for AMR profile of transformants [87]. Antimicrobials tested were as follows (μg/disk): erythromycin (15), clindamycin (2), gentamicin (10), kanamycin (30), streptomycin (10 U), tobramycin (10), tetracycline (30), trimethoprim (5), sulfonamide (300) and TMP–sulfamethoxazole (1.25 + 23.75).

#### Phenotypic characterization of biofilm formation

Biofilm formation ability of the four *S. sciuri* strains as well as *S. aureus* RN4220/pUR2865-34 transformants was tested by a modified Congo Red Agar (CRAmod) assay and by Cristal Violet microtitre plate assay (S1 Data for detailed procedures) [88, 89]. *S. aureus* SA113 and *S. aureus* ATCC 25923 were used as positive controls for their strong biofilm-forming potential, while *S. aureus* RN4220 (DSM 26309) was selected as negative control for biofilm production.

#### PCR detection of the TMP resistance gene *dfrE* and the *ica* locus variant

Primers designed for the detection of the TMP resistance *dfrE* gene, the biofilm formation *ica* locus (*ica*ADBC) variant genes and the *ica* locus repressor (*icaR*) are indicated in S1 Table. For the *dfrE* gene, primers designed enclosed the Dhfr superfamily domain region (Cd-Search PF00186) and all *dhfr* genes with ≥ 45% nucleotide similarity were included to search for specificity of *dfrE* gene primer set. Original strains (C2853, C2854, C2855), RN4220/pUR2865-34 transformants and several control strains were tested for sensitivity and specificity (S1 Data).

## Acknowledgements

The authors would like to thank Prof. Carmen Torres, from the University of La Rioja (Spain), and Prof. Kennedy Chah, from the University of Nigeria Nsukka, for kindly providing us with *S. sciuri* strains C2865, C2853, C2854 and C2855, and Dr. Brion Duffy and Dr. Theo Smits, from the Zurich University of Applied Sciences (ZHAW), Wädenswil (Switzerland), for the Illumina sequencing raw data of *S. sciuri* C2865. We would also like to thank Prof. Vincent Perreten and Dr. Sybille Schwendener, from the Institute of Veterinary Bacteriology, Bern (Switzerland) for the vectors pBUS-HC and pBUS-Pcap-HC, as well as Prof. Francisco Rodriguez-Valera for accessibility to computing resources. We thank Dominique Lorgé for excellent technical assistance. We would also like to thank Prof. Gloria del Solar, Prof. Jose R. Penadés and Dr. Alexandro Varani for expert discussion on plasmid and pathogenicity island biology, respectively, and bioinformatic analysis support related to ISs. Part of these data was presented at the Swiss General Meeting (SGM) of the Swiss Society for Microbiology (SSM) Joint Annual Meeting in Basel (August 30 – September 01, 2017), Switzerland; in the American Society for Microbiology (ASM) 2nd ASM Conference on Rapid Applied Microbial Next-Generation sequencing and Bioinformatic Pipelines (October 8-11, 2017), Washington DC, USA; and in the 5th International Symposium on the Environmental Dimension of Antibiotic Resistance (EDAR-5) (June 9 – 14, 2019) in Hong Kong.

## Author contribution statements

**Conceptualization:** Elena Gómez-Sanz

**Data curation:** Elena Gómez-Sanz, Jose M. Haro-Moreno, Mario López-Pérez, Slade O. Jensen

**Formal analysis:** Elena Gómez-Sanz, Mario López-Pérez, Jose M. Haro-Moreno

**Funding acquisition:** Elena Gómez-Sanz

**Investigation:** Elena Gómez-Sanz

**Methodology:** Elena Gómez-Sanz, Jose M. Haro-Moreno, Mario López-Pérez

**Project administration:** Elena Gómez-Sanz

**Resources:** Elena Gómez-Sanz + see acknowledgements

**Software:** Jose M. Haro-Moreno, Mario López-Pérez

**Supervision:** Elena Gómez-Sanz

**Validation:** Elena Gómez-Sanz, Mario López-Pérez, Jose M. Haro-Moreno, Juan José Roda-García, Slade O. Jensen, Francisco Rodriguez-Valera

**Visualization:** Elena Gómez-Sanz, Mario López-Pérez, Jose M. Haro-Moreno

**Writing ± Original Draft Preparation:** Elena Gómez-Sanz

**Writing – Review & Editing:** Elena Gómez-Sanz, Mario López-Pérez, Jose M. Haro-Moreno, Slade O. Jensen

## Data availability statement

The whole genome shotgun sequence data supporting this work are available via de NCBI SRA database under Bioproject and Biosample accession numbers PRJNA663854 and SAMN16182282, respectively. Individual accession numbers are as follow: xxx (S. sciuri C2865 chromosome with PacBio RSII), xxx (plasmid pUR2865-1), xxx (plasmid pUR2865-2), and xxx (plasmid pUR2865-34).

## Supporting information

**S1 Data. Supplementary data for detailed material and methods.**

**S1 Figure. Dot-plot graphs of MRSS C2865 genome assembled by PacBio and by Illumina uding D-Genies.** The program searches all the query sequences aligned on the diagonal and is calculating the target sequence coverage per identity bin [72]. Left graph shows the genome-genome alignments using PacBio assembly as reference (chromosomal contig and plasmid contig) (x axes) with Illumina query (y axes), while right graph depicts the genome-genome alignments using Illumina assembled contigs (n=as reference (x axes) with PacBio query (y axes). The summary identity calculation (>75% identity, <75%, <50% and <25% and no matches) is made on the target sequence (reference) and represents the percentage of the target genome in base pairs. Note: Node 96 (5’513 bp) denotes plausible contamination after contig manual checking.

**S2 Figure. Phylogenomic tree and heat map resultant from the Average Nucleotide Identity (ANI) of 30 Staphylococcus sciuri group genomes.** S. sciuri genomes included are S. sciuri NS36 (GenBank assembly accession: GCA_001477335), *S. sciuri* NS1 (GCA_001476955), *S. sciuri* NS44 (GCA_001477405), *S. sciuri* RSA37 (GCA_001476585), *S. sciuri* NS53 (GCA_001476575), *S. sciuri* NS112 (GCA_001477395), *S. sciuri* ATCC29059 (GCA_900117375), *S. sciuri* FDAARGOS285 (GCA_002209165.), *S. sciuri* LCHXa (GCA_002091355), *S. sciuri* NCTC12103 (GCA_900474615), *S. sciuri* DSM20345 (GCA_001046995), *S. sciuri* MC10 S56 (GCA_003006205), *S. sciuri* P575 (GCA_001766775), *S. sciuri* S P879 (GCA_001766785), *S. sciuri* SAP15-1 (GCA_001684285), *S. sciuri* NS202 (GCA_001476555), *S. sciuri* i1 (GCA_002407445), *S. sciuri* SNUD18 (GCA_002072755), *S. sciuri* CCUG39509 (GCA_002902225), *S. sciuri* Z8 (GCA_000612145), *S. sciuri* C2865 (Biosample acc. No. SAMN16182282), *S. lentus* MF1767 (GCA_001651345), *S. lentus* 050AP (GCA_900098655), *S. lentus* MF1862 (GCA_001651255), *S. lentus* NCTC12102 (GCA_900458735), *S. lentus* F1142 (GCA_000286395), *S. vitulinus* DSM15615 (GCA_002902265), *S. vitulinus* F1028 (GCA_000286335), *S. fleuretti* MBTS1 (GCA_002018435), *S. stepanovicii* NCTC13839 (GCA_900187075) plus one *Staphylococcus aureus* and one *Staphylococcus epidermidis* reference genomes (*S. aureus* NCTC8325 [GenBank RefSeq. Acc. No. NC_007795] and *S. epidermidis* ATCC12228 [NC_004461]), used as outgroup for the geneus level.

**S3 Figure. Neighbor-joining tree of aligned amino acid sequences of *S. sciuri* group dihyfrofolate reductases.** The alignment was performed with (i) the entire dihydrofolate reductase (Dhfr) protein/s present in all *S. sciuri* species group genomes deposited in the NCBI database (as of April 2018), (ii) both Dhfrs present in *S. sciuri* strain C2865, (iii) the amino-acid sequence of all trimethoprim resistance dihydrofolate reductases (Dfr) described so far in staphylococci (DfrA, DfrD, DfrG, DfrK, DfrF) [35-39], (iv) as well as the Dhfr of trimethoprim susceptible *S. epidermidis* ATCC12228 (NC_004461) and *S. aureus* ATCC25923 (Z16422) as reference. Of note, when more than one *ddfr* gene was present per genome, both resultant proteins were included and labelled as “Dhfr1” and “Dhfr2”, based on the genome position as deposited in NCBI, respectively. Amino acid sequence alignment was performed using Muscle, with UPGMB as clustering method. The tree was built by the Neighbour-joining method with a bootstrap of 1000 replications (substitution model: poison model; rates among sites: Gamm Distribution). Gene product, staphylococcal species and strain are followed by their GenBank accession numbers within brackets. The three Dhfr present in *S. sciuri* C2865 are displayed in bold. The amino-acid branch clustering the trimethoprim resistance DfrE is highlighted in faint green, which also encloses the trimethoprim resistance DfrF. Subcluster enclosing additional staphylococcal trimethoprim resistance Dfrs (DfrD, dfrG and DfrK) have a faint blue background. The remaining trimethoprim resistance DfrA is clustered with both trimethoprim susceptible Dfrs selected as reference (blue font) and the sub-branch is emphasised on faint yellow background.

**S4 Figure. Biofilm formation ability of original *ica*-locus variant carrying *S. sciuri* strains and transformants.** (A) Biofilm formation capacity based on visual analysis of colony colours by a modified Congo Red Agar (CRAmod) assay. Spots were plated in duplicates. From left to right, 1) displays the positive controls (strong biofilm formers) *S. aureus* SA113 (DSM 4910) and *S. aureus* ATCC 25923 (DSM 1104); 2) negative control (pUR2865-34 recipient) strain *S. aureus* RN4220; 3) shows the original *S. sciuri ica*-locus variant carrying strains C2865, C2853, C2854 and C2855; 4) includes two selected *S. aureus* RN4220/pUR2865-34 transformants (S319 and S320). (B) Quantification of biofilm formation by crystal violet (CV) staining of adherent cells. The bar chart displays the results of six independent experiments in triplicates. Original *S. sciuri ica*-locus variant carrying strains C2865, C2853, C2854 and C2855, two selected *S. aureus* RN4220/pUR2865-34 transformants (S319 and S320) as well as an *ica*-negative non biofilm producing methicillin-resistant *Staphylococcus lentus* strain (C3030) [107] are included. The three bars on the left show the control strains used: the strong biofilm formers *S. aureus* SA113 (DSM 4910) and *S. aureus* ATCC 25923 (DSM 1104), as well as *S. aureus* RN4220 strain. Significant differences by t-test or ANOVA (P < 0.01) are indicated with two stars on the compared cluster, i.e. *ica*-carrying strains (C2865, C2853, C2854, C2855, S319, S320; ns, not significant differences between values) and C3030; and *ica*-carrying strains and RN4220. Strains producing a mean absorbance value of > 0.3 were considered weak biofilm producers, while those with >1.0 strong biofilm producers.

**S5 Figure. Graphical comparative analysis of pUR2865-1 (A) and pUR2865-2 (B) with the closest sequences in NCBI.** (A) Represented closest plasmids similar to pUR2865-1 (min 80% ID, min 70% coverage) and strains carrying them are as follow (GenBank acc. no): *S*. *aureus* strain C5425 plasmid pUR5425, complete sequence (JQ861958.1); *S. aureus* strain SR434 plasmid pSR04, complete genome (CP019567.1); *Staphylococcus chromogenes* plasmid pLNU4, isolate KNS48 (NC_007771.1); *Staphylococcus simulans* plasmid pLNU2, isolate 184/61 (NC_007769.1). (B) Represented plasmids closest to pUR2865-2 (min 80% ID, min 70% coverage) and strains containing them are as follow (GenBank acc. no): *S*taphylococcus hyicus strain 1211 plasmid pSWS1211 (KM276081.1); Staphylococcus aureus plasmid pSWS2889, strain MRSA ST398, isolate 1110902889 (NC_023385.1); S. aureus plasmid pDJ91S, complete sequencec (KC895984.1); S. aureus chloramphenicol resistance plasmid pKH7, complete sequence (U38429.1); S. aureus plasmid pOC160-2 DNA, complete sequence, strain: OC160 (LC012933.1); and *S. aureus* R-plasmid pSBK203 replication initiation protein gene, chloramphenicol acetyltransferase gene, and Pre protein gene, complete cds (U35036.1).

**S6 Figure. Graphical overview of MRSS C2865 ψTn*554* chromosomal region and comparative elements.** The diagram depicts (i) MRSS C2865 ψTn*554* chromosomal region including the interrupted *radC* gene, (ii) its corresponding closest relative *S. aureus* 6850 (Genbank acc. No. CP006706), and (iii) the chromosomal *radC* region and remnant 3’-end *radC* of phenicol and oxazolidinone resistant *S. sciuri* wo22_7 (KX982170) and *S. sciuri* MS11-3 (KX447571), respectively. Arrows denote the genes, length and orientation. Gene colours other than dark turquoise represent the following genes of interest (colour): truncated or remnant chromosomal integration radC gene in Tn554 family transposons (black); antimicrobial resistance genes (pink); genes involve in metal resistance or transport (bright blue); genes involved in transposition or recombination (yellow); and genes coding for ABC transporter ATP-binding proteins (faint brown). The 6-bp nucleotides resultant from Tn*554*-like integration are also depicted. Areas of nucleotide similarity (nblastn, >100 bp match, >85% ID) between strains/structures are indicated in grey.

**S7 Figure. Comparison of codon usage (A) and amino acid (B) percentage between the three prophages (C2865-pp1, C2865-pp2, C2865-pp3) and C2865 host genome.** The scale (in percentage) in the left plots is represented at the upper center of the graph. (A) Left, radar p lot of the 64 different amino acid codons (at DNA level) highlighting that of tRNA gene present in prophage C2865-pp3 genome. Right, specific abundance (%) of TGG codon (UGG). (B) Left, radar plot of the 20 amino acids highlighting proline (P), as resultant amino acid from tRNA present in C2865-pp3. Right, specific abundance (%) of proline (P).

**S8 Figure. Phylogenomic tree and heat map resultant from the Average Nucleotide Identity (ANI) of 187 staphylococcal phages deposited in the Viral RefSeq NCBI data base.** The upper left-hand site displays the clusterization of all 190 phages included, while the main heat map represents the phylogenomic grouping of phages within the sub-branch enclosing the three novel prophages only (C2865-pp1, C2865-pp2 and C2865-pp3).

**S9 Figure. Circular phylogenetic tree of all available staphylococcal terminase large subunit (TerL).** Circular phylogenetic tree built with the amino acid sequences of all identifiable terminase large subunit (TerL) present in 187 staphylococcal phages deposited in the Viral RefSeq database, TerL of C2865-pp1, C2865-pp2 and C2865-pp3, and TerL of Enterobacteria phage lambda (RefSeq, acc. No. NC_001416), used as reference. Phage nomenclature: S-phage, which refers to Staphylococcal phage, is followed by the phage name and. faconcant plus the CDSs number of respective TerL in the phage genome. Amino acid sequence alignment was performed using CLUSTALW and the tree was built by the Neighbour-joining method with a bootstrap of 500 replications (protein substitution model: WAG) (CLC Genomics Workbench 11). Branch enclosing the C2865-pp1, C2865-pp2 and C2865-pp3 integrases is bold. C2865-pp1, C2865-pp2 and C2865-pp3 integrases are boxed in green, while phage lambda is boxed in grey. Phages with known packaging mechanism [60], carrying either *cos*-sites (*cos* packaging strategy) or *pac*-sites (headful packaging strategy), are indicated in black and green, respectively.

**S1 Table.**
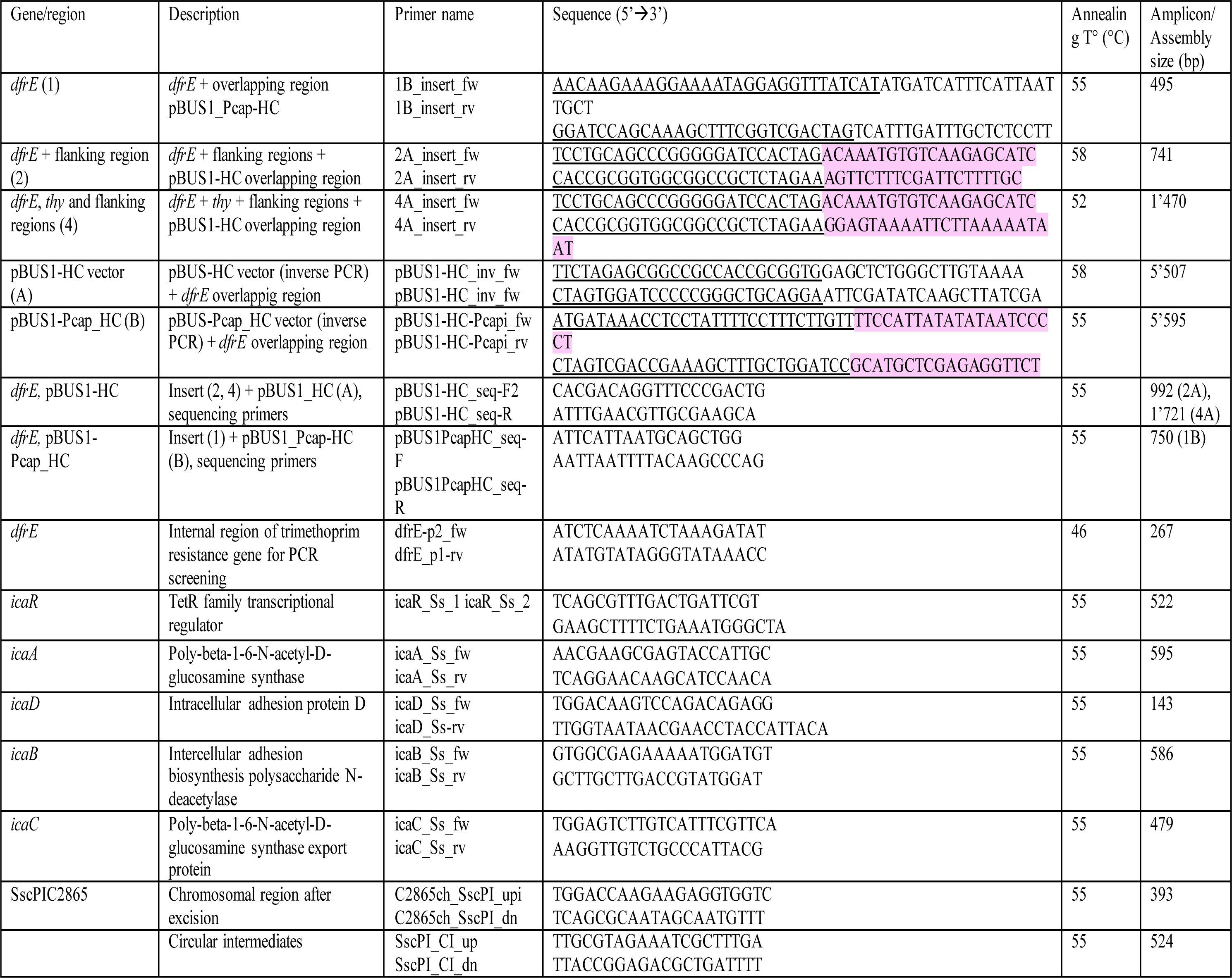
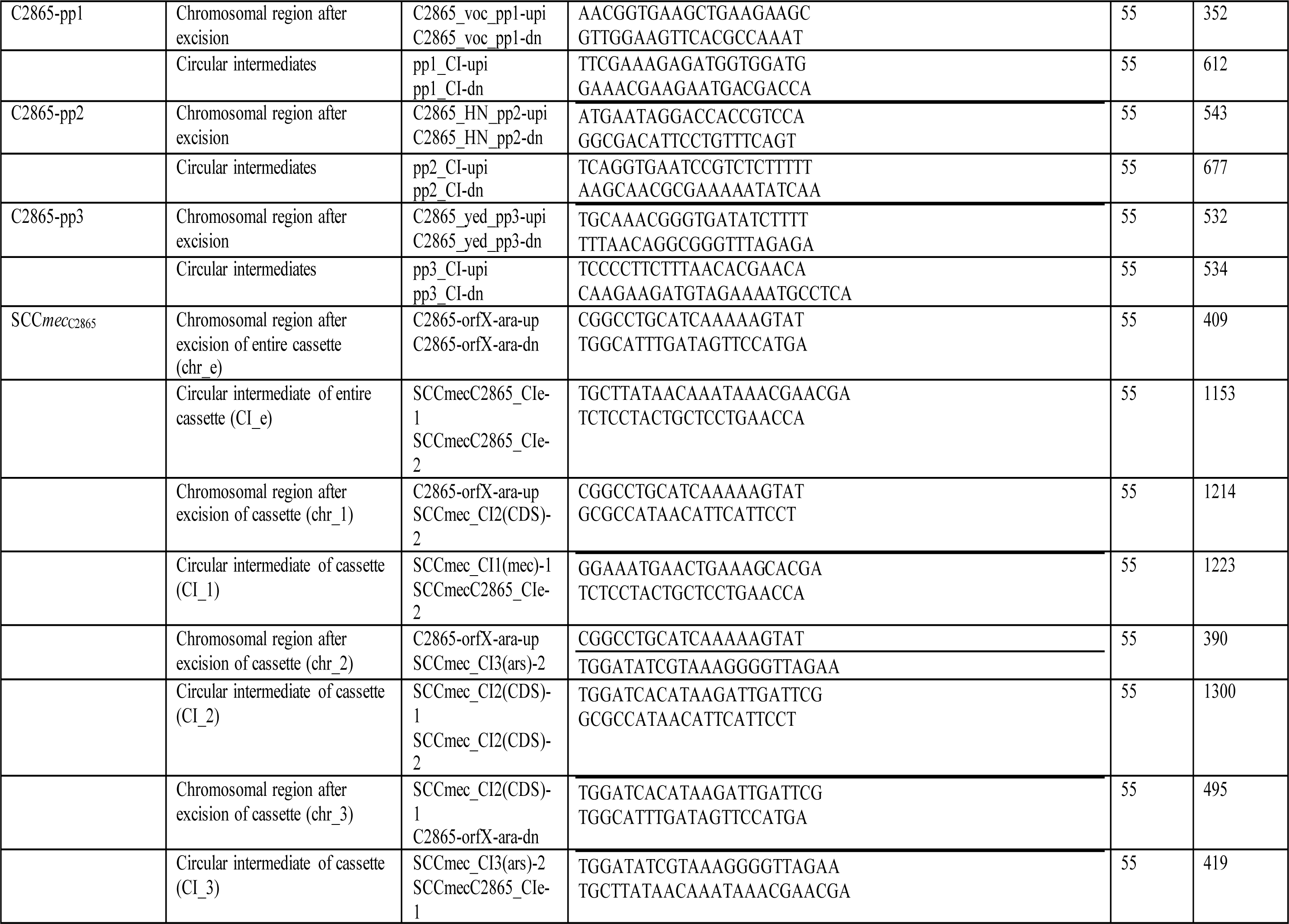
Primers designed and used in this study.

